# Morpho-physiological and transcriptomic responses of field pennycress to waterlogging

**DOI:** 10.1101/2024.08.15.608142

**Authors:** Rachel Combs-Giroir, Manesh Shah, Hari Chhetri, Mallory Morgan, Erica Prates, Alice Townsend, Mary E. Phippen, Winthrop B. Phippen, Daniel A. Jacobson, Andrea R. Gschwend

**Affiliations:** Center for Applied Plant Sciences, The Ohio State University, Columbus, OH, USA; Department of Horticulture and Crop Sciences, The Ohio State University, Columbus, OH, USA; Biosciences, Oak Ridge National Laboratory, Oak Ridge, TN, USA; The Bredesen Center for Interdisciplinary and Graduate Education, University of Tennessee, Knoxville, TN, USA; School of Agriculture, Western Illinois University, Macomb, IL, USA

**Keywords:** *Thlaspi arvense*, transcriptomics, waterlogging, hypoxia, RNA-seq, recovery, root stress, ERF-VII

## Abstract

Field pennycress (*Thlaspi arvense*) is a new biofuel winter annual crop with extreme cold hardiness and a short life cycle, enabling off-season integration into corn and soybean rotations across the Midwest. Pennycress fields are susceptible to winter snow melt and spring rainfall, leading to waterlogged soils. The objective of this research was to determine if waterlogging during the reproductive stage had a significant effect on gene expression, morphology, physiology, recovery, and yield of two pennycress lines (SP32-10 and MN106). In a controlled environment, total pod number, shoot/root dry weight, and total seed count/weight were significantly reduced in SP32-10 in response to waterlogging, whereas primary branch number, shoot dry weight, and single seed weight were significantly reduced in MN106. This indicated waterlogging had a greater negative impact on seed yield in SP32-10 than MN106. The number of differentially expressed genes (DEGs) between waterlogged and control roots were doubled in MN106 (3,424) compared to SP32-10 (1,767). Functional enrichment analysis of upregulated DEGs revealed Gene Ontology (GO) terms associated with hypoxia and decreased oxygen, with genes in these categories involved in alcoholic fermentation and glycolysis. Interestingly, MN106 waterlogged roots exhibited significant stronger upregulation of these genes than SP32-10. Additionally, downregulated DEGs revealed GO terms associated with cell wall biogenesis and secondary metabolite biogenesis, indicating suppressed growth and energy conservation. Together, these results reveal the reconfiguration of cellular and metabolic processes in response to the severe energy crisis invoked by waterlogging in pennycress.

## Introduction

Flooding is a serious threat to global agriculture, causing drastic economic losses from reduced crop production and quality (Rosenzweig et al., 2002; Mirza, 2011). Global warming has led to the increased frequency of heavy precipitation and flooding events. Studies using global climate modeling data project precipitation increases up to 30% in the spring and winter months across the Midwest and Great Lakes regions in the 2080s, resulting in increased soil moisture (Demaria et al., 2016; Byun and Hamlet, 2018; Byun et al., 2019), and statistical modeling predicts continued increases in flood events throughout the 21^st^ century (Mallakpour and Villarini, 2015; Neri et al., 2020). This poses the risk of crop damage within the U.S. corn belt region, particularly for crops growing during the spring months, such as field pennycress.

Waterlogging occurs when the root zone of a plant becomes saturated or flooded, leading to a low-oxygen, or hypoxic, environment where the diffusion of oxygen and other gases is restricted (Bailey-Serres et al., 2012). This inhibits oxidative phosphorylation, leading to low levels of ATP production. In most plants, waterlogging typically results in reduced stomatal conductance, root hydraulic conductivity, and nutrient and water uptake (Striker, 2012; Liu et al., 2020), as well as an increase in toxic metabolites and reactive oxygen species (ROS) (Perata and Alpi, 1991; Gibbs and Greenway, 2003; Pucciariello and Perata, 2017). These effects can lead to chlorophyll degradation, leaf chlorosis, and early senescence, resulting in reduced photosynthetic rate and eventually plant death (Ashraf and Mehmood, 1990; Ahmed et al., 2012; Liu et al., 2012; Striker, 2012; Xu et al., 2019). To combat the severe energy shortage under hypoxia, plants will upregulate glycolysis, fermentation, and sugar metabolic processes (Sairam et al., 2008; van Dongen and Licausi, 2015).

Field pennycress (*Thlaspi arvense* L.) is a winter annual oilseed crop native to Eurasia and distributed across temperate regions in North America (Sedbrook et al., 2014). Pennycress is a promising bioenergy crop for the U.S. Midwest Corn Belt due to its high seed oil content and fatty acid oil composition that meets the U.S. renewable fuels standards when converted to biofuel (Moser et al., 2009; Moser, 2012). The extreme cold hardiness and short life cycle of pennycress allow off-season integration into corn and soybean rotations, preventing displacement of summer commodity crops (Fan et al., 2013; Sedbrook et al., 2014). As a winter annual crop, pennycress has potential to prevent nutrient leaching, soil erosion, and spring weed growth, while also providing a food source for native pollinators (Eberle et al., 2015; Johnson et al., 2017; Weyers et al., 2019). These economic and ecological benefits have led to the domestication of pennycress for improved yield, agronomic traits, and oil/protein quality, resulting in a cash crop that has the potential to produce up to 1 billion liters of seed oil annually by 2030 (Phippen et al., 2022). Pennycress is typically planted in autumn (September-October), flowers in early April, and matures in mid to late May meaning that pennycress plants are typically in the reproductive stages when heavy spring precipitation events occur. Anecdotal evidence suggests pennycress is susceptible to damage in areas of fields with standing water (communication with Win Phippen), though this has not been documented in controlled experimental investigations to date. For pennycress to become a successful cash cover crop, improved resiliency to abiotic threats faced during the growing season will be essential.

Pennycress research is bolstered by a vast molecular toolkit consisting of a reference genome and transcriptome for the MN106 winter annual line (Dorn et al., 2013, 2015; Nunn et al., 2022), an EMS mutant gene index (Chopra et al., 2018), a sequenced genome of an inbred spring annual line called Spring 32-10 (McGinn et al., 2019), and optimized protocols for transformation and gene editing (McGinn et al., 2019). Pennycress is grouped into lineage 2 of the *Brassicaceae* family with *Brassica* spp. and is closely related to lineage 1 plants, such as *Arabidopsis thaliana* and *Camelina sativa* (Best and Mcintyre, 1975; Dorn et al., 2013). Thus, the abundant data and resources available for *Arabidopsis* and related oilseed crops are easily translatable to pennycress, especially due to a general one-to-one correspondence between *Arabidopsis* genes and pennycress orthologs (Chopra et al., 2018).

Waterlogging and submergence stress have been well studied in the model crop *A. thaliana* and *Brassica napus* (rapeseed), but is lacking for other *Brassicaceae* crops such as *Brassica rapa, Brassica oleraceae*, *Camelina sativa*, and pennycress (Combs-Giroir and Gschwend, 2024). Furthermore, transcriptomic studies of *Brassicaceae* crops under waterlogging stress at reproductive stages are limited. This work seeks to bridge these gaps by first characterizing the morphological and physiological responses of pennycress to waterlogging at the reproductive stage, and second, investigating the transcriptomic responses of pennycress root and shoot tissue under waterlogging. Our results revealed differences in morphology and yield under waterlogging stress between the two evaluated lines, as well as molecular differences, therefore, providing candidate genes and pathways involved in differential responses to waterlogging. This data can be further utilized by breeders and scientists to improve pennycress resiliency to heavy spring precipitation events.

## Materials and methods

### Plant Material and Growth Conditions

The pennycress accession MN106 (winter-type) originates from Coates, Minnesota (ABRC stock: CS29149) and is the source of the reference genome, and SP32-10 (spring-type) originates from Bozeman, Montana (ABRC stock: CS29065) and is a rapid-cycling, sequenced lab line. MN106 and SP32-10 seeds were sterilized with a 70% ethanol rinse, followed by 10-minute incubation in 30% bleach/0.01% SDS solution, and then rinsed 3 times with sterile water. Seeds were then plated on petri dishes between two pieces of Whatman paper with 2mL sterile water, stratified at 4°C for 5 days, and moved to a growth chamber at 21°C with periods of 16-hour light/8-hour dark. Following 1 week of germination, the MN106 seedlings were vernalized at 4°C under 14hrs of daily light for 3 weeks to induce flowering. Approximately 2 weeks after germination or vernalization, the seedlings were transplanted into plug cells in Berger BM2 Germination Mix (Hummert International, Earth City, MO, United States) and then into 10-cm pots after the first true leaves fully developed. The plants were grown at ∼21°C with periods of 16-hour light/8-hour dark either in a growth chamber or a greenhouse. The media of the pots was 50% Turface MVP calcined clay (PROFILE Products LLC, Buffalo Grove, IL United States), 25% PRO-MIX BX MYCORRHIZAE (Premier Horticulture, Quakertown, PA, United States), and 25% Miracle-Gro All Purpose Potting mix (Scotts Miracle-Gro, Marysville, OH, United States). Plants were watered with Jack’s Nutrients 12-4-16 RO (jacksnutrients.com) at 100ppm. The water was dechlorinated with 2.5mg/L of sodium thiosulfate pentahydrate early in plant development after transplanting.

### Greenhouse Experimental Design

Following approximately 2 weeks of flowering, during pennycress developmental stage 6 (Verhoff et al., 2022), 18 plants for each accession were placed in 7.6-liter buckets and waterlogged for one week by filling water up to the soil line at the top of the pots. An additional 18 plants for each accession were also placed in buckets but watered normally approximately every other day so that the soil was completely saturated after watering, but dry before the next watering. Our experimental design was per the following: six blocks per accession, each block containing two buckets (one waterlogged and one control), each bucket containing three plants of the same accession. The buckets for each accession were randomized on a greenhouse bench in a random complete block design. Since MN106 plants flowered about 1 week later than SP32-10, the waterlogging treatments for each accession took place a week apart. Morphological data was collected immediately after waterlogging where 6 plants per treatment were destructively harvested and measured for plant height, reproductive height (total height-height to first silicle), primary branch number, silicle number, total leaf number, and shoot and root fresh and dry weight. Root and shoot tissues were oven-dried at 50°C for 48 hours. Additionally, 12 plants per treatment/control were allowed to recover and were measured at maturity, when the plants had fully senesced, for plant height, reproductive plant height, silicle number, percentage of aborted silicles, the average number of seeds per silicle for 10 silicles, branch number (total and primary), days until maturity, total dry shoot and root weight, 1000 seed weight, total seed count, total seed weight, and single seed weight.

### Growth Chamber Experimental Design

The greenhouse experiment described above was repeated in a growth chamber using the same growth conditions, in an even ratio of the three soil types, but not watered with Jack’s nutrients. The same experimental design was implemented, but there were only 12 plants/replicates for each treatment. Immediately after waterlogging, 6 plants per treatment were destructively harvested for RNA-seq (see below) and the other 6 plants were measured for morphological traits 1) immediately after waterlogging, 2) after 1 week of recovery, 3) after 2 weeks of recovery, and 4) at maturity when the plants were harvested. The following phenotypes were collected: inflorescence status, silicle number, percentage of aborted and senesced silicles, plant height, reproductive plant height, branch number (total and primary), days until maturity, total dry shoot weight, 1000 seed weight, total seed count, total seed weight, and single seed weight.

### Total Seed Oil Content

Total oil content was determined by nondestructive TD-NMR on 450 mg samples of whole pennycress seed. TD-NMR was performed on the samples and standards with a Bruker Minispec MQ40 with a 39.95 MHz NMR frequency and 40°C magnet temperature. The 90° and 180° pulse lengths were 11.48 µs and 22.98 µs respectively. A 23° detection angle, gain of 44 dB, pulse attenuation of 15 dB, recycle delay of 2 s, window 1 of 0.055 ms, window 2 (τ_ε_) of 7 ms, and 724 magnetic field steps were set to employ a Hahn spin echo effect for signal collection. Each sample and standard analysis employed 16 scans at a receiver gain of 59. The percent oil for each sample was calculated using the Bruker Oil and Moisture in Seeds application in minispec Plus version 7.0.0 software.

A percent oil calibration curve was constructed within the Bruker software by measuring the signal for six oil standards between 20% −40% oil concentration based on a 450 mg sample. Crude pennycress oil in six amounts ranging from 90 mg to 180 mg was carefully dropped down six 10 mm flat bottom NMR tubes containing a piece of kimwipe in the bottom. After the oil standards were tempered at 40°C for 30 minutes, they were scanned in the MQ40 in the calibration mode to create a signal vs concentration calibration curve for percent oil.

To determine percent oil in seeds from the sample plots, an analytical balance was used to weigh 450 mg of seed from each plot into 10 mm flat bottom NMR tubes. After tempering the samples at 40°C for 30 minutes, they were scanned in the MQ40 and the percent oil was calculated using our calibration curves and the Bruker Oil and Moisture in Seeds application in minispec Plus software. The results from minispec Plus were exported to an Excel file. Using Excel, the percent oil on a dry weight basis was calculated by dividing the milligrams of oil in each sample by the dry weight of each sample and multiplying by 100.

### Fatty Acid Seed Oil Composition

For gas chromatography (GC) analysis of oil components, medium-chain triglycerides were extracted and converted to fatty acid methyl esters (FAMEs) through a transesterification reaction. A Mettler Toledo analytical balance was used to weigh 100 mg of seed from each plot into 20 mL glass scintillation vials. To each sample, 5mL of 0.25 M sodium methoxide in methanol was added, and an Ingenieurburo CAT X 120 homogenizer was used to thoroughly grind the seed. The vials were capped and placed in an oven at 60°C. After 30 minutes, the vials were removed and cooled to room temperature. To each vial, 2 mL of saturated sodium chloride in distilled water and 5mL of hexane were added. The vials were shaken and two separate layers developed. Approximately 1.5 mL of the top layer which was comprised of hexane with FAMEs was pipetted into a GC autosampler vial for GC analysis.

The fatty acid methyl esters were analyzed using an Agilent 6890 gas chromatograph with a 7683 autosampler and flame ionization detector (FID). A Supelco 2380 30m x 0.25mm x 0.2µm film thickness column was used. Ultra-high purity helium was the carrier gas with a constant pressure of 20.00 psi. A 1 µL injection was used with a split ratio of 50:1. Split flow and total flow were 58.3 mL/min and 62.2 mL/min. For the detector, ultra-high purity hydrogen and air were used in the FID with flow rates of 40.0 mL/min and 450.0 mL/min respectively. The injector and detector temperatures were set at 265°C and 250°C respectively. The oven temperature was set initially at 170°C and ramped at 4°C/min to 190°C then ramped at 30°C/min to 265°C and held for 2.5 min. NuChek standard 17A and Supelco 37 component FAME standard were run to match retention times. Data was collected and the area percents for the FAMEs were exported into an Excel file using Agilent OpenLab CDS ChemStation Edition revision C.01.10 software.

### Physiological Data Collection

During the greenhouse experiment, stomatal conductance (gsw), CO_2_ assimilation (A), and transpiration rate (E) were collected on 12 plants per treatment per accession 1 day before waterlogging, at 7 days of waterlogging, and at 7 days of recovery during the greenhouse experiment using the LI-6800 Portable Photosynthesis System (Licor, Lincoln, NE, USA). Measurements were collected between ∼10am and 12pm with a healthy leaf on the main stem. The instrument flow rate was set to 400 μmol/s, relative humidity at 50%, CO_2__s at 400 ppm, fan speed at 10,000 rpm, and the light source at ambient light conditions. The instrument was stabilized on A, gsw, and fluorescence prior to logging a measurement.

## Statistics

Normality and homogeneity of variance were confirmed using Shapiro-Wilks and Levene’s tests. Traits that did not have equal variances for one or both accessions were plant height, reproductive plant height, shoot fresh weight, percentage of aborted silicles, and thousand seed weight. A Welch’s t-test was used to determine significance between waterlogged and control replicates of each trait for both accessions waterlogged at the reproductive stage. A two-way analysis of variance (ANOVA) followed by a post-hoc Tukey’s Honestly Significant Difference (HSD) test with alpha = 0.05 was used to determine the effects of treatment, accession, and their interaction. For physiological traits, a repeated measures ANOVA was used with time as a within-subject factor and accession and treatment as between-subject factors to determine the effects of timepoint, accession, treatment, and their interaction. Values for gsw, A, and E were log transformed. Statistically significant main effects were followed by pairwise t-tests with Bonferroni correction for multiple comparisons. All statistical analyses were conducted in RStudio (R version 4.2.1).

### Tissue Collection and RNA isolation

Leaves, silicles, and roots were collected from 5 separate plants (replicates) per accession from the growth chamber experiment after 1 week of waterlogging and after 3 hours of recovery. Tissues from control plants were also collected after 1 week, coinciding with the collection of the waterlogged plants. Additionally, leaves, silicles, and roots were collected from SP32-10 waterlogged and control plants after 3 days of waterlogging. Roots were gently washed in water for ∼2 minutes to remove soil and Turface. Tissues were flash-frozen in liquid nitrogen and kept at −80°C until RNA isolation. RNA was isolated from the leaves and roots using the Plant RNeasy Qiagen kit (qiagen.com), and silicles were isolated using the Spectrum Plant Total RNA Kit (sigmaaldrich.com) following kit protocols. RNA samples were stored at −80°C until sent for mRNA library preparation and paired-end Illumina sequencing (NovaSeq 6000 PE150) at Novogene Corporation Inc.

### RNA-seq Analysis

Raw sequence read quality was analyzed using FASTQC v0.11.8 (Babraham Bioinformatics - FastQC A Quality Control tool for High Throughput Sequence Data, n.d.) before and after trimming low-quality reads and adaptor sequences using Trimmomatic v0.38 (Bolger et al., 2014). One sample (MN106 control roots – replicate 1) was removed from the analysis due to poor sequence quality. Clean sequence reads were aligned to the version 2 pennycress genome (GenBank GCA_911865555.2) using STAR v2.7.9a (Dobin et al., 2013). Raw read counts were assigned using featurecounts v2.0.1 (Liao et al., 2014). MultiQC v1.13 was used after FASTQC, STAR, and featurecounts to aggregate output files and obtain summary of results (Ewels et al., 2016). All processing steps were performed with the Ohio Supercomputer Center (Ohio Supercomputer Center., 1987).

Raw read counts were tested for intersample differential expression (DE) analysis using the DESeq2 v1.38.3 (Love et al., 2014) and “apeglm” to shrink the log2FoldChange (Zhu et al., 2019) in RStudio v4.2.1. The negative binomial GLM was fitted and Wald statistics were calculated, followed by identification of differentially expressed genes using the Benjamini-Hochberg method with a false discovery rate (FDR) < 0.05 and a relative change threshold of 2-fold or greater. A test for interaction of accession and condition was used to reveal genes significantly different between accessions for a given condition or between time points. Volcano plots were generated using the EnhancedVolcano R package v1.16.0 (Blighe et al., 2023). Heatmaps were generated using the pheatmap R package v1.0.12 (Kolde, 2019).

### GO and KEGG Enrichment Analysis

Pennycress genes were assigned an *A. thaliana* gene ID based on shared homology using OrthoFinder v2.5.4 (Emms and Kelly, 2019). Gene Ontology (GO) and Kyoto Encyclopedia of Genes and Genomes (KEGG) enrichment of the RNA-seq results was carried out with the Gene Set Enrichment Analysis (GSEA) method using the clusterProfiler R package v4.6.2 and the *A. thaliana* database (Wu et al., 2021) with a p-value cutoff < 0.05. Shrunken LFC values were used as input.

### MENTOR algorithm

We identified 2,089 unique significantly differentially expressed genes (DEGs) for control vs waterlogged MN106 samples and 432 unique DEGs from control vs waterlogging SP32-10 samples (Log_2_FC = ± 1.3). We combined the DEGs for both varieties and used the MENTOR (Multiplex Embedding of Network Topology for Omics Resources) algorithm (Sullivan et al., 2024) to group functionally or mechanistically connected genes into clusters unique to each genotype. In brief, the input for MENTOR is a set of genes that are the seed for a Random Walk with Restart (RWR) algorithm starting from each gene and traversing the network to find functionally (or mechanistically) connected genes based on the network topology. Here, *Arabidopsis thaliana* genes orthologous to *Thlaspi arvense* DEGs were used as starting genes. Random walks traversed an *Arabidopsis thaliana* multiplex network composed of literature curated and exascale/petascale generated gene sets derived from experimental data (Kainer et al., 2024; Sullivan et al., 2024). The output of MENTOR is a dendrogram with clusters of genes ordered by their functional or mechanistic relationships to one another, which was determined by their connectivity among the genes in the multiplex network (Sullivan et al., 2024). A heatmap with up- and downregulated genes for MN106 and SP32-10 in waterlogged relative to control plants is also associated with this dendrogram.

## Results

### Morphology and physiology immediately after waterlogging

To determine if waterlogging affects pennycress growth, we waterlogged MN106 and SP32-10 accessions for 7 days and recorded morphology and physiology data immediately after waterlogging. These experiments were carried out in a greenhouse and repeated in a growth chamber. The only significant changes to growth and reproduction between control and waterlogged plants immediately following the treatment was a significant reduction (∼32%) in leaf number for MN106 waterlogged plants compared to controls (Figure 1, Table 1, and Supplemental Table 1) and a significantly higher percentage of dead inflorescences (11%) in SP32-10 waterlogged plants compared to the controls (0%) (Supplemental Figure 1 and Supplemental Table 2). Furthermore, root dry weight was reduced by 26% in MN106 and 19% in SP32-10, and shoot dry weight by 9.3% in MN106 and 2.7% in SP32-10, though not significantly (Table 1). Transpiration rate, stomatal conductance, and CO_2_ assimilation were significantly reduced in both accessions after 7d of waterlogging (Supplemental Figure 2A-C).

**Figure 1.**
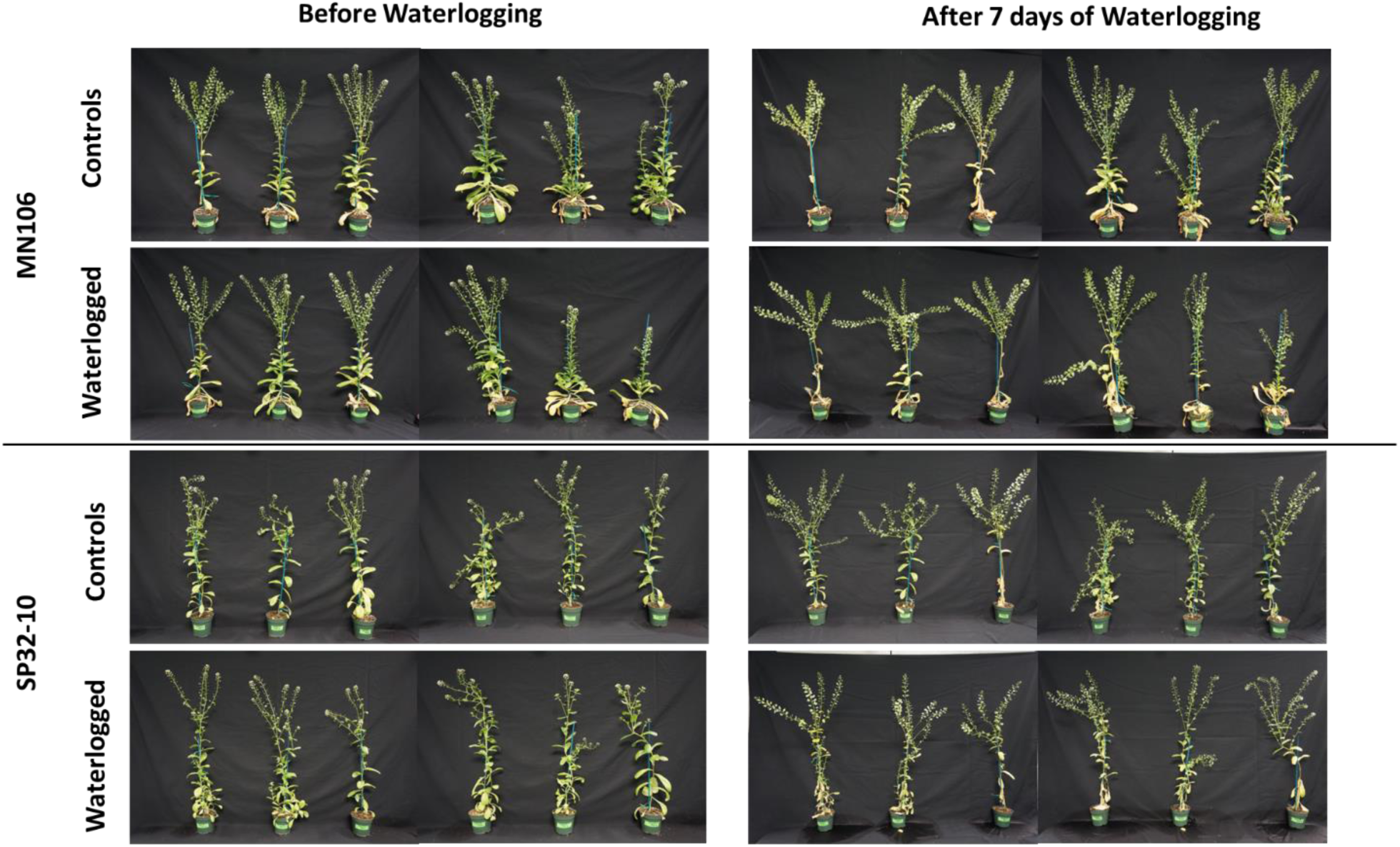
Reproductive-stage pennycress plants before and after 7 days of waterlogging in the greenhouse experiment.

**Table 1.**
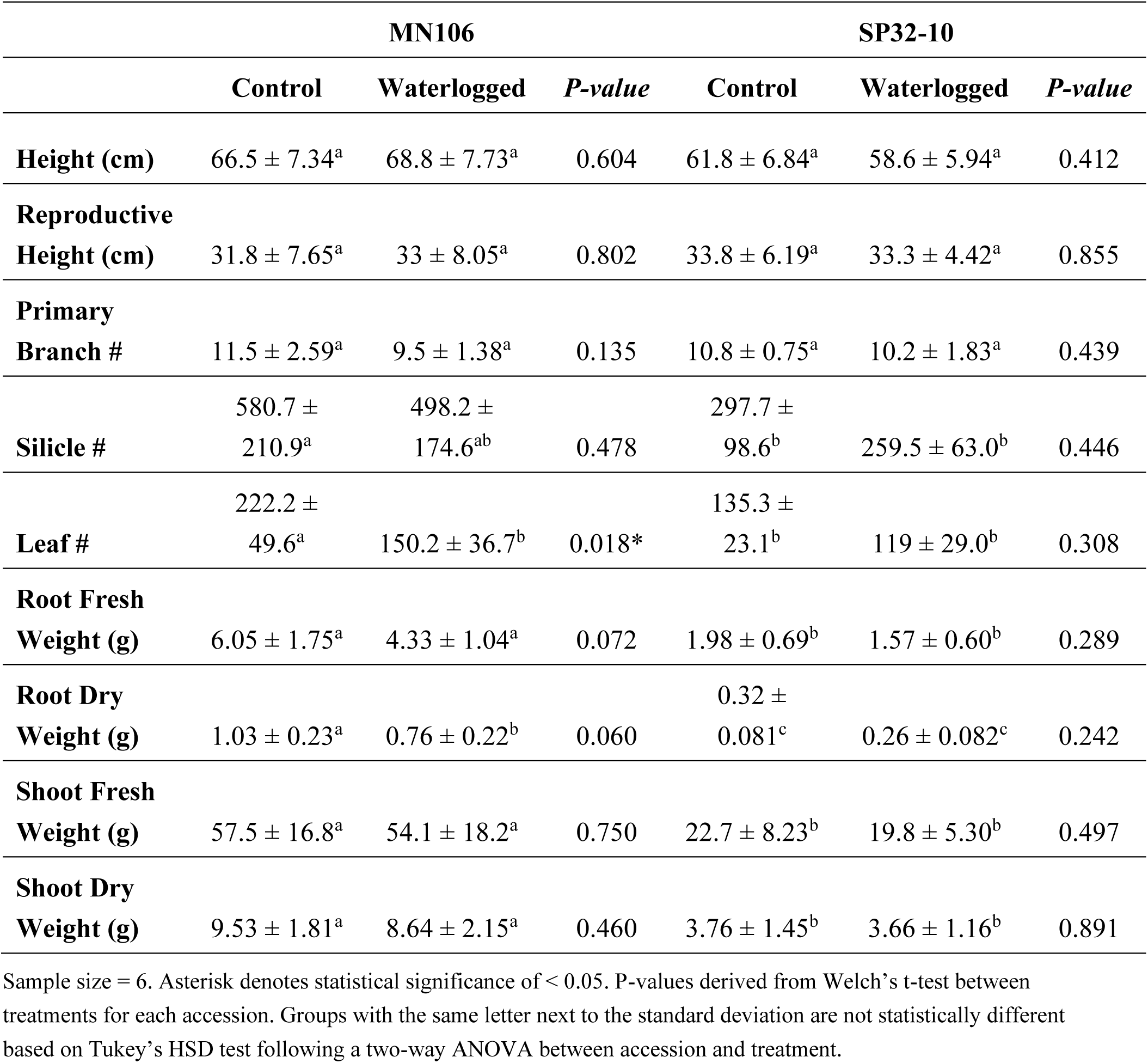
Means and standard deviations of morphological traits of waterlogged and control plants immediately after waterlogging in the greenhouse experiment.

### Morphology and physiology during recovery

To determine whether waterlogging had lasting effects on pennycress growth and development, plants were observed during the first two weeks of recovery. SP32-10 waterlogged plants had significantly more senesced silicles than controls (Supplemental Figure 3, Supplemental Table 3). Transpiration rate and CO_2_ assimilation remained significantly reduced in SP32-10 after 1 week of recovery (Supplemental Figure 2A-C).

### Morphology, biomass, and yield at the time of harvest

Morphological traits, biomass, and yield were recorded once the plants had fully matured and senesced (Table 2). In SP32-10, waterlogged plants had a significant reduction in the total number of silicles, with an average of about 713 silicles compared to an average of 1,138 silicles in control plants. MN106waterlogged plants had a significant decrease in mean total primary branches, 11 compared to 16 total primary branches in control plants, but this reduction did not significantly reduce silicle number. Additionally, shoot dry weight of waterlogged plants was significantly reduced by 23.7% in MN106 and 27.3% in SP32-10. The total root dry weight of waterlogged plants was significantly reduced by 41% in SP32-10 and reduced by 30% in MN106, but not significantly. The percentage of aborted silicles was significantly increased by 25% in SP32-10 after waterlogging in the repeated growth chamber experiment (Supplemental Table 4).

**Table 2.** Means and standard deviations of morphological traits of waterlogged and control plants at the time of harvest in the greenhouse experiment.

|  | MN106 |  |  | SP32-10 |  |  |
| --- | --- | --- | --- | --- | --- | --- |
|  | Control | Waterlogged | *P-value* | Control | Waterlogged | *P-value* |
| **Height (cm)** | 65.4 ± 6.93<sup>a</sup> | 64.2 ± 8.46<sup>a</sup> | 0.908 | 66.7 ± 2.93<sup>a</sup> | 66.3 ± 5.34<sup>a</sup> | 0.833 |
| **Reproductive Height (cm)** | 29.1 ± 7.19<sup>a</sup> | 29.6 ± 9.38<sup>a</sup> | 0.875 | 36.1 ± 4.63<sup>a</sup> | 34.8 ± 3.31<sup>a</sup> | 0.460 |
| **Primary Branch #** | 15.8 ± 4.85<sup>a</sup> | 11.4 ± 3.18<sup>b</sup> | 0.018* | 10.6 ± 2.61<sup>b</sup> | 10.9 ± 3.50<sup>b</sup> | 0.794 |
| **Branch #** | 59 ± 37.45<sup>ab</sup> | 36.7 ± 23.22<sup>b</sup> | 0.096 | 82.4 ± 22.35<sup>a</sup> | 65.8 ± 34.60<sup>ab</sup> | 0.179 |
| **Silicle #** | 911.5 ± 472.5<sup>ab</sup> | 623.1 ± 232.8<sup>b</sup> | 0.076 | 1138.4 ± 416.8<sup>a</sup> | 712.9 ± 318.4<sup>b</sup> | 0.011* |
| **Aborted Silicles (%)** | 42.5 ± 18.8<sup>a</sup> | 34.5 ± 27.6<sup>a</sup> | 0.198 | 35.4 ± 11.9<sup>a</sup> | 27.9 ± 8.78<sup>a</sup> | 0.095 |
| **Maturity** | 48.3 ± 9.22<sup>ab</sup> | 45.3 ± 11.1<sup>b</sup> | 0.931 | 59.1 ± 4.5<sup>a</sup> | 51.9 ± 10.8<sup>ab</sup> | 0.019* |
| **Root Dry Weight** | 0.60 ± 0.32<sup>ab</sup> | 0.42 ± 0.13<sup>ab</sup> | 0.347 | 0.61 ± 0.14<sup>a</sup> | 0.36 ± 0.16<sup>b</sup> | 0.001*** |
| **Shoot Dry Weight** | 15.6 ± 4.26<sup>a</sup> | 11.9 ± 4.24<sup>ab</sup> | 0.046* | 13.9 ± 2.69<sup>ab</sup> | 10.1 ± 3.97<sup>b</sup> | 0.013* |
| **Total Seed Count** | 3235.2 ± 1011.2<sup>a</sup> | 2906.1 ± 1142.4<sup>a</sup> | 0.462 | 4546.7 ± 1007.1<sup>a</sup> | 3148.9 ± 1148.1<sup>a</sup> | 0.003** |
| **Total Seed Weight (g)** | 3.73 ± 1.45<sup>ab</sup> | 3.0 ± 1.18<sup>b</sup> | 0.186 | 4.61 ± 1.05<sup>a</sup> | 3.16 ± 1.45<sup>b</sup> | 0.011* |
| **Thousand Seed Weight (g)** | 1.15 ± 0.27<sup>a</sup> | 1.06 ± 0.39<sup>a</sup> | 0.266 | 1.01 ± 0.05<sup>a</sup> | 0.93 ± 0.12<sup>a</sup> | 0.143 |
| **Single Seed Weight (mg)** | 0.93 ± 0.093<sup>ab</sup> | 0.78 ± 0.18<sup>b</sup> | 0.019* | 1.11 ± 0.15<sup>a</sup> | 1.00 ± 0.35<sup>ab</sup> | 0.344 |
| **Seeds per**<br>**silicle** | 8.79 ±<br>1.05<sup>a</sup> | 8.72 ± 0.76<sup>a</sup> | 0.844 | 9.43 ± 0.76<sup>a</sup> | 8.74 ± 0.83<sup>a</sup> | 0.045* |
| **Oil Content**<br>**( % DWB)** | 30.1 ±<br>2.19<sup>a</sup> | 28.1 ± 2.13<sup>ab</sup> | 0.120 | 27.7 ±<br>2.46<sup>bc</sup> | 26.8 ± 3.70<sup>c</sup> | 0.360 |

Sample size = 12. Asterisk denotes statistical significance of < 0.05. DWB = dry weight basis. P-values derived from Welch's t-test between treatments for each accession. Groups with the same letter next to the standard deviation are not statistically different based on Tukey's HSD test following a two-way ANOVA between accession and treatment.

Several parameters of seed yield were also measured (Table 2 and Supplemental Figure 4). Compared to controls, SP32-10 waterlogged plants had a significant reduction in total seed count by 31%, total seed weight by 31%, and seeds per silicle by 7%. The repeated growth chamber experiment also presented similar results (Supplemental Table 4 and Supplemental Figure 5). Waterlogged MN106 plants only had a significant reduction in single seed weight by 16% compared to controls (Table 2 and Supplemental Figure 4D), which was also observed in the repeated growth chamber experiment (24% reduction) (Supplemental Table 4 and Supplemental Figure 5D).

Since pennycress is harvested for seed oil, we wanted to test if waterlogging caused any changes in oil content or fatty acid profiles in the mature seed. In MN106, there was no significant difference in total oil content after waterlogging in the greenhouse experiment (Table 2 and Supplemental Figure 4F), but there was a significant 10.6% reduction in the repeated growth chamber experiment (Supplemental Table 4 and Supplemental Figure 5E). In SP32-10, no significant differences were detected in total oil content. MN106 waterlogged seeds showed several significant changes in fatty acid constituents compared to control seeds following the greenhouse experiment, such as an increase in palmitic acid and linoleic acid and decreases in eicosenoic acid, erucic acid, and constituents grouped into the “other” category (Figure 2A). Only a significant increase in behenic acid in MN106 was observed in the repeated growth chamber experiment (Supplemental Figure 6). SP32-10 waterlogged seeds only showed a significant decrease in oleic acid (Figure 2B).

**Figure 2.**
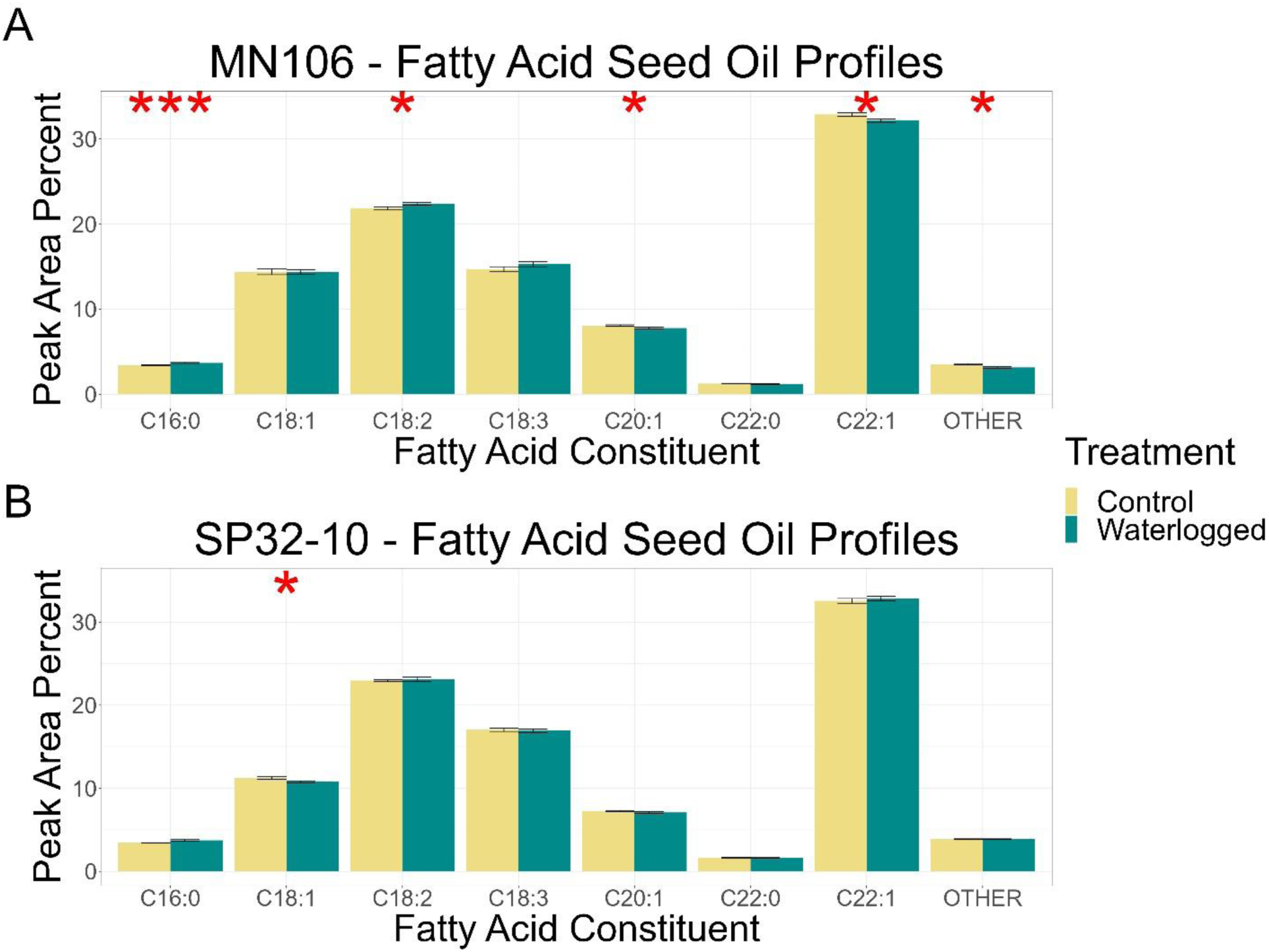
Fatty acid oil profiles of mature seed after waterlogging treatment in the greenhouse experiment in (**A**) MN106 and (**B**) SP32-10. Fatty acid constituents are C16:0 = palmitic acid, C18:1 = oleic acid, C18:2 = linoleic acid, C18:3 = linolenic acid, C20:1 = eicosenoic acid, C22:0 = behenic acid, C22:1 = erucic acid. * = p-value < 0.05, ** = p-value < 0.01, ***=p-value < 0.001, bars denote standard error.

### Differentially expressed genes between waterlogged and control plants (MN-7WL and SP-7WL)

To determine the genes contributing to a response to waterlogging stress, we waterlogged plants for 7 days following 2 weeks of flowering and compared the transcriptome of the roots, leaves, and silicles to plants watered normally for both MN106 (MN-7WL) and SP32-10 (SP-7WL). After 7 days of waterlogging, despite few phenotypic changes in the plants, there was a large transcriptomic response. A total of 3,424 genes (∼13% of genes in the genome) were differentially expressed in MN-7WL roots, with 1,590 genes upregulated and 1,834 downregulated, whereas only 1,767 genes (∼7% of total genes) were differentially expressed in SP-7WL roots, with 974 genes upregulated and 793 downregulated under waterlogging (Figure 3A, Supplemental Figure 7, Supplemental File 1). Significantly fewer differentially expressed genes (DEGs) were identified between the waterlogged and control pennycress leaves and silicles (Supplemental Figure 7). After 7 days of waterlogging, there were only 27 DEGs, 26 upregulated and 1 downregulated, in MN106 leaves, whereas no DEGs were detected in the leaves of SP32-10. There were 37 and 2 DEGs in MN106 and SP32-10 silicles, respectively, all upregulated compared to the controls. These results suggest MN106 had a broader transcriptome response to waterlogging (more DEGs in more tissues) compared to SP32-10 and roots had the greatest transcriptomic response to waterlogging compared to shoot tissues directly following waterlogging. Therefore, we moved forward with analyses on the root transcriptome data.

**Figure 3.**
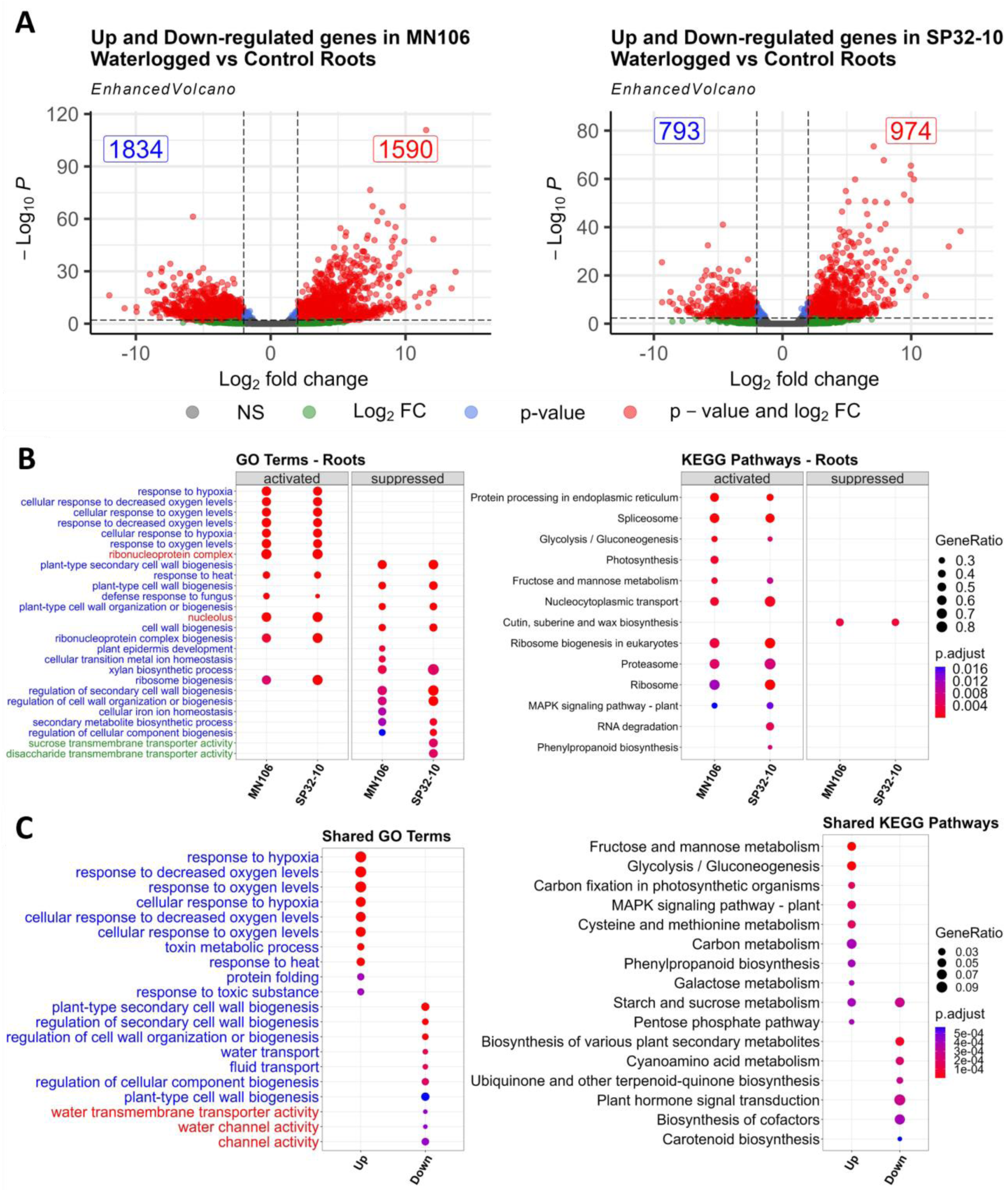
Differentially expressed genes and enrichment results for waterlogged roots compared to controls. (**A**) Volcano plots showing the log2FC of downregulated and upregulated differentially expressed genes (red dots) in waterlogged roots based on cutoffs of |log2FC| ≥ 1 and a p-value ≤ 0.05, (**B**) Dot plot of 10-30 most significant gene ontology and KEGG terms of gene set enrichment analysis in MN-7WL and SP-7WL roots. Blue, red, and green terms represent biological process, cellular component, and molecular function, respectively. GeneRatio represents the ratio of the number of genes from the input data that belong to each term out of the total number of genes in the database associated with that term and p.adjust represents the level of significance of the enrichment. (**C**) Over-representation analysis of 10-30 most significant gene ontology and KEGG terms from the shared overlapping up- and downregulated DEGs between MN-7WL and SP-7WL.

Next, we performed a gene set enrichment analysis (GSEA), which ranks all genes by the log2 fold change (log2FC) value to look at the entire gene landscape, to see which GO terms and KEGG pathways were enriched in MN-7WL and SP-7WL roots. We found 128 enriched GO terms in MN-7WL and 135 in SP-7WL where 100 had positive enrichment scores (genes were mostly upregulated) and about 30 terms had negative enrichment scores (genes were mostly downregulated) (Supplemental File 2, Figure 3B). Genes involved in 12 KEGG pathways were enriched in MN-7WL and 14 in SP-7WL, and only 1-2 pathways had downregulated genes. In MN-7WL and SP-7WL roots, upregulated genes were significantly enriched for GO terms related to ribonucleoprotein complex, response to hypoxia/decreased oxygen, and response to heat, whereas downregulated genes were significantly enriched for GO terms related to cell wall biogenesis, xylan biosynthetic process, and secondary metabolite biosynthetic process. Out of 199 genes in the database belonging to response to hypoxia, 115 (58%) upregulated genes contributed to the enrichment for MN-7WL and 105 (53%) for SP-7WL.Additionally, out of the 118 genes in the database for plant-type secondary cell wall biogenesis, 63 (53%) downregulated genes contributed to the enrichment for MN-7WL and 72 (61%) for SP-7WL. This indicates that more than half of the hypoxia and secondary cell wall biogenesis-related genes were differentially expressed in our dataset. We then looked at the significant DEGs within these categories to see their respective functions (Supplemental File 3). Some of the upregulated DEGs under waterlogging in the hypoxia category were *PRYUVATE DECARBOXYLASE 2 (PDC2)*, *ALCOHOL DEHYDROGENASE 1 (ADH1)*, *SUCROSE SYNTHASE 1 (SUS1)*, *HYPOXIA RESPONSIVE ERFs (HRE1* and *HRE2*). The top 10 most significant hits can be found in Supplemental Table 5. Furthermore, some significant DEGs in the secondary cell wall biogenesis category had roles in lignin and cellulose biosynthesis and cell morphology, such as *LACCASE 10 (LAC10)*, *MYB DOMAIN PROTEIN 20 AND 58 (MYB20*, *MYB58)*, *C-TERMINALLY ENCODED PEPTIDE RECEPTOR 1* (*CEPR1*), and *CELLULOSE SYNTHASE A4* (*CESA4*). Additionally, upregulated genes were significantly enriched for KEGG pathways related to the spliceosome/ribosome/proteasome, nucleocytoplasmic transport, fructose and mannose metabolism, and glycolysis/gluconeogenesis, whereas downregulated genes were significantly enriched for cutin, suberin, and wax biosynthesis (Supplemental File 4, Figure 3B). Out of the 96 glycolysis genes, 30 genes were enriched in both accessions, and 12 out of 26 cutin/suberine/wax biosynthesis genes were enriched in both accessions. Significant DEGs in the glycolysis category were mostly genes encoding glycolytic enzymes, such as *FRUCTOSE-BISPHOSPHATE ALDOLASEs (FBA6*, *FBA3)*, *HEXOKINASE 3 (HXK3)*, *ENOLASE 2 (ENO2)*, *PHOSPHOENOLPYRUVATE CARBOXYKINASE 2* (*PCK2)*, while significant DEGs involved in cutin/suberin/wax biosynthesis consisted of *FATTY ACID REDUCTASEs (FAR5*, *FAR4), CYTOCHROME P450 (CYP86)*, *ABERRANT INDUCTION OF TYPE THREE 1* (*ATT1)*, and *CALEOSIN 6 (CLO6)*, which have roles in suberin biosynthesis and fatty acid metabolic processes (Supplemental File 3). These enrichment results, and the significant DEGs within the enriched categories, indicate changes in gene expression related to stress responses, metabolism, transcription/translation, and development. Furthermore, several of the top 10 upregulated DEGs in waterlogged roots for each accession (Supplemental Table 5) are involved in ethylene synthesis and stress responses like fermentation, such as *WOUND-INDUCED POLYPEPTIDE 4 (WIP4)*, *PDC2*, *ADH1*, *GLUTAMATE DECARBOXYLASE 4* (*GAD4)*, *ACC OXIDASE 1 (ACO1)*, and *LOB DOMAIN-CONTAINING PROTEIN 40 (LBD40)*, further supporting some of the stress response gene enrichment categories detected with the GSEA analyses.

We also performed an over-representation analysis (ORA) of the genes that were differentially expressed in both MN-7WL and SP-7WL roots (Figure 3C). A total of 1,335 genes (768 upregulated, 566 downregulated, and 1 gene with contrasting expression) were differentially expressed in the roots of both accessions (Figure 4A). Shared upregulated genes were enriched in GO categories such as response to hypoxia, response to heat, toxin metabolic process, and protein folding, whereas shared downregulated genes were enriched in categories such as secondary cell wall biogenesis, water transport, and water channel activity. Genes contributing to the water transport enrichment were primarily aquaporin genes, such as *PLASMA MEMBRANE INTRINSIC PROTEINS* (*PIP2;2, PIP1, PIP1;4, PIP1;5, PIP2;6*), *ATP-BINDING CASSETTE G22*, and *TONOPLAST INTRINSIC PROTEIN 2;3*, with log2FC ranging from −2.4 to −5.9. Additionally, shared upregulated genes were enriched in KEGG pathways such as fructose and mannose metabolism, glycolysis, carbon fixation and metabolism, and MAPK signaling, whereas shared downregulated genes were enriched in categories such as plant secondary metabolite biosynthesis, cyanoamino acid metabolism, and plant hormone signal transduction.

**Figure 4.**
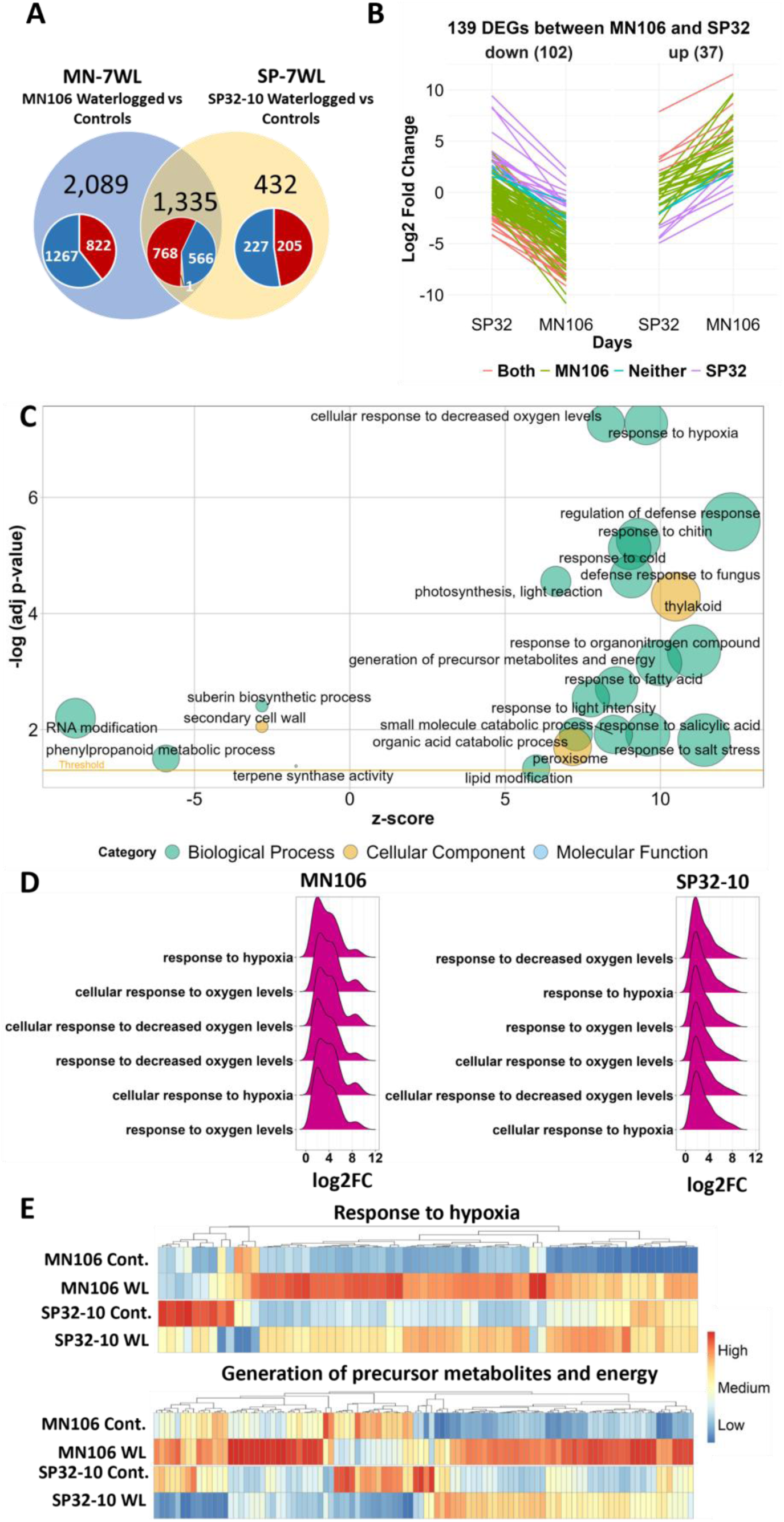
Differentially expressed genes (DEGs) and enrichment results between accessions in waterlogged compared to control roots. (**A**) Venn diagram of overlapping DEGs from MN106 waterlogged vs controls (MN-7WL) and SP32-10 waterlogged vs controls (SP-7WL). Red, blue, and orange colors in pie chart represent upregulated, downregulated, and contrasting gene expression respectively. (**B**) The change in log2FC values of the 102 down and 37 upregulated DEGs in the interaction from MN-7WL vs SP-7WL. Coral-colored lines indicate genes that were significant in both MN-7WL and SP-7WL, green and purple lines indicate genes significant in only MN-7WL and SP-7WL respectively, and blue lines indicate genes that were not significant in either MN-7WL or SP-7WL. (**C**) Bubble plot of GO terms from GSEA results of the interaction from MN-7WL and SP-7WL. The size of the bubbles are proportional to the number of genes assigned to the term and the z-score provides an estimate if the genes are more likely to be upregulated or downregulated and is calculated by (# of upregulated genes - # of downregulated genes)/√count. (**D**) Ridge plots showing log2FC values of genes falling under hypoxia-related enriched GO terms in the interaction between MN-7WL and SP-7WL. The height of the peak corresponds to the number of genes with the respective log2FC value. (**E**) Heatmap of genes across treatments and accessions falling under the ‘response to hypoxia’ and ‘generation of precursor metabolites and energy’ GO terms from GSEA interaction analysis between MN-7WL and SP-7WL. Heatmap scale was applied by column or gene.

### Differentially expressed genes between accessions after waterlogging (MN-7WL vs SP-7WL)

Since there were differences in the total number of DEGs between MN106 and SP32-10 in response to 7d waterlogging, we further investigated which of these DEGs were uniquely differentially expressed in the roots between the accessions. A total of 2,089 genes (61% of total MN-7WL root DEGs) were uniquely differentially expressed in MN-7WL and 432 genes (25% of total SP-7WL root DEGs) were uniquely expressed in SP-7WL (Figure 4A). To find the genes differentially regulated between the two accessions in response to waterlogging, the interaction, or change in log2FC, between the treatment and the accessions was calculated (MN-7WL vs SP-7WL). This comparison revealed 37 upregulated and 102 downregulated DEGs in MN-7WL compared to SP-7WL (Figure 4B, Supplemental File 5). Out of these 139 DEGs, 95 were also significant in, and specific to, MN-7WL, whereas 18 were significant in, and specific to, SP-7WL. The remaining 26 DEGs were either significant in both MN-7WL and SP-7WL (10 DEGs) or were oppositely expressed between MN-7WL and SP-7WL (16 DEGs). For instance, TAV2_LOCUS6692, a gene encoding a carotenoid isomerase, was upregulated by a log2FC of about 3 in MN-7WL but downregulated by a log2FC of about 2 in SP-7WL. While this gene was not detected as significantly differentially expressed between waterlogged and control roots in either SP32-10 or MN106, it was identified as having a significant interaction between accessions, because of the opposite expression pattern. Among the 95 DEGs specific to MN-7WL, 71 were downregulated and 24 were upregulated. For the 18 DEGs specific to SP-7WL, 13 were upregulated and 5 were downregulated.

Next, we ranked all the genes by the change in the log2FC and employed a GSEA to look at the entire gene landscape. This revealed 39 positively enriched GO terms, such as response to oxygen levels/hypoxia, regulation of defense response, response to chitin, response to cold, photosynthesis and thylakoid, generation of precursor metabolites and energy, response to fatty acid, peroxisome, and others (Figure 4C and Supplemental File 2). The function of the DEGs in MN-7WL vs SP-7WL also overlapped with these GSEA results. For instance, some of the 37 upregulated DEGs in MN-7WL vs SP-7WL are known to respond to abscisic acid, hypoxia, starvation, and water deprivation such as *CYTOSOLIC ABA RECEPTOR KINASE 5 (CARK5,* Δlog2FC = 2.3), *3-KETOACYL-COA SYNTHASE 4* (*KCS4,* Δlog2FC = 2.3), *PHYTOCYSTATIN 4 (CYS4,* Δlog2FC = 2.6), an O-Glycosyl hydrolases family 17 protein (Δlog2FC = 2.9), *COLD-RESPONSIVE 6.6* (*COR6.6,* Δlog2FC = 3.2), *XYLOGLUCAN ENDOTRANSGLUCOSYLASE/HYDROLASE 11* (*XTH11,* Δlog2FC = 3.2), *ATPS2* (Δlog2FC = 5.5), and a 2-oxoglutarate (2OG) and Fe(II)-dependent oxygenase superfamily protein (Δlog2FC = 10.6). Interestingly, *ATPS2* was among the top 10 most significant DEGs in MN-7WL with a log2FC of 8.7 (Supplemental Table 5). Additionally, upregulated unique genes (no Arabidopsis ortholog), based on homology to proteins in other species, are involved in cell wall modification such as *PHOSPHOLIPASE D Z* (Δlog2FC = 3.8), *PECTATE LYASE* (Δlog2FC = 5.4), *PECTINESTERASE* (Δlog2FC = 8.7), and *XTH* (Δlog2FC = 2.5).

On the other hand, downregulated genes were enriched for 6 GO terms in the GSEA, such as RNA modification, suberin biosynthetic process, secondary cell wall, casparian strip, phenylpropanoid metabolic process, and terpene synthase activity (Figure 4C and Supplemental File 2). Genes that contributed to the core enrichment of the secondary cell wall and casparian strip GO categories consisted of *ENHANCED SUBERIN 1* (*ESB1*, TAV2_LOCUS14251), *LACCASE 1* (*LAC1*, TAV2_LOCUS1633), *MYB DOMAIN PROTEIN 39* (*MYB39,* TAV2_LOCUS24544), and *CASPARIAN STRIP MEMBRANE DOMAIN PROTEIN 3* (*CASP3*/TAV2_LOCUS14279) which were all significantly downregulated in MN-7WL by a log2FC of about 3, except for *CASP3* which was downregulated by a log2FC of about 7 (Supplemental File 1). None of these genes were differentially expressed in SP-7WL, and the latter three were significant DEGs in MN-7WL vs SP-7WL (Δlog2FC = −4 - 5.8) (Supplemental File 5). Some of the 102 downregulated DEGs were also involved in cell wall loosening, lignan biosynthesis, cell growth, and casparian strip lignification, such as *EXPANSIN 18* (*EXP18,* Δlog2FC = −6.7), *PINORESINOL REDUCTASE 2* (*PRR2,* Δlog2FC = −5.1), *PEROXIDASE 9* (*PER9*, Δlog2FC = −5) and *PEROXIDASE 39* (*PER39,* Δlog2FC = −4.6), and two CAP superfamily proteins (Δlog2FC = −7). Many of the unique pennycress genes in the MN-7WL vs SP-7WL DEG list also point at the capability of MN106 to break down cell wall components under root waterlogging. For instance, the downregulated unique pennycress genes in MN-7WL vs SP-7WL were *EXTENSION-2 like* genes (Δlog2FC = −6.4), *INDOLE GLUCOSINOLATE O-METHYLTRANSFERASES* (Δlog2FC = −5.5 and −4.4), *WALL-ASSOCIATED RECEPTOR KINASES* (Δlog2FC = −4.2), *CASP* (Δlog2FC = −6.1), and a germin-like protein (Δlog2FC = −6.5). Lastly, some of the downregulated DEGs with the highest fold changes in MN106 waterlogged roots compared to SP32-10 are involved in iron and sulfur starvation and root development, such as *BASIC HELIX-LOOP-HELIX PROTEIN 100* (*BHLH100,* Δlog2FC = −10.2), *IRON-REGULATED TRANSPORTER 1* (*IRT1,* Δlog2FC = −9.4), *NICOTIANAMINE SYNTHASES* (*NAS*, Δlog2FC = −9 and −7.1), a Pollen Ole e 1 allergen and extension family protein (Δlog2FC = −6.9), and *UDP-GLUCOSYL TRANSFERASE 76C4* (*UGT76C4,* Δlog2FC = −6.2).

While many of the 139 DEGs had similar functions to the GSEA GO categories, not all of them were found in the enrichment results, indicating that these GO categories contained groups of genes that collectively had smaller changes in expression levels between the two accessions. For instance, MN106 waterlogged roots had stronger expression of genes involved in the response to hypoxia than SP32-10 with a log2FC of about 4 in MN106 and about 2 in SP32-10 (Figure 4D&E). Some of the genes contributing to the core enrichment of this category were *SENESCENCE ASSOCIATED GENE 14 (SAG14*) and *RESPIRATORY BURST OXIDASE HOMOLOG D (RBOHD)*, which doubled in expression in MN106 waterlogged roots compared to SP32-10. MN106 waterlogged roots also had stronger expression of genes involved in the generation of precursor metabolites and energy (Figure 4C). Some of the genes contributing to the core enrichment of this category consisted of *PYRUVATE KINASE FAMILY PROTEINS* and *HXK-like 1*, which were significantly upregulated in MN-7WL but not in SP-7WL. Additionally, *GACPC2*, *FBA6,* and *ENO2* were also significantly upregulated in waterlogged roots of both accessions but were higher in MN-7WL by a log2FC of about 0.5-1 (Supplemental File 1).

To explore functionally connected genes unique to each accession, we used the MENTOR algorithm to assign genes into clusters, which are represented in a dendrogram (Supplemental Figure 8). This included 913 genes that were differentially expressed in the roots of one genotype but not the other. We used the associated heatmaps to identify large clusters of functionally related genes that were significantly upregulated or downregulated in each genotype to mechanistically interpret genotype-specific responses to waterlogging. Examination of the functional patterns in these clades indicated that many of them involved ion transporters and efflux pumps, osmotic stress, hormone regulation and response, and lipid biosynthesis with several connections to suberin. We selected one extended cluster for deeper interpretation. In the context of waterlogging stress, the regulation and function of specific pennycress genes exemplified in this functional clade (all but one down-regulated under waterlogging stress) highlight differences in adaptive strategies between MN106 and SP32-10 genotypes, to cope with the challenges posed by excessive water and reduced oxygen availability.

The genes in this clade support a narrative of an intricate balance of water and ion homeostasis, essential for plant survival during waterlogging. For instance, we observed the downregulation of several genes in MN-7WL involved in ion balance and osmotic stability, such as *KUP6* (TAV2_LOCUS15590/AT1G70300), *KAT3/AtKC1* (TAV2_LOCUS22773/AT4G32650), *NIP5;1* (TAV2_LOCUS19334/AT4G10380), and *PHO1;H1* (TAV2_LOCUS16940/AT1G68740). The only upregulated gene in this functional clade was in MN-7WL and encodes a MATE Efflux Family Protein (TAV2_LOCUS1072/AT1G11670), which could be a response to prevent the accumulation of toxic metabolites in the cells or to maintain hormonal homeostasis under abiotic stress. Furthermore, MN-7WL had several downregulated genes involved in root hydrotropism and meristem growth, such as a gene encoding the MIZU-KUSSEI-like (MIZ) protein (TAV2_LOCUS5684/AT5G23100), *BAM3* (TAV2_LOCUS24776/AT4G20270), and *SKU5 Similar 15* (TAV2_LOCUS24681/AT4G37160).

Additionally, there were several downregulated genes involved in suberin and lipid metabolism in MN-7WL, such as a gene encoding an HXXXD-type acyl-transferase family protein (TAV2_LOCUS2538/AT1G24430) and *GGL25* (TAV2_LOCUS19216/AT5G03610). Lastly, we identified several downregulated genes in MN-7WL implicated in hormone responses such as *DIOXYGENASE FOR AUXIN OXIDATION 1* (*DAO1*/ TAV2_LOCUS1366/AT1G14130) and *RESPONSIVE-TO-ANTAGONIST 1* (*RAN1*/ TAV2_LOCUS6093/AT5G44790), as well as those previously mentioned such as *KUP6*, *PHO1;H1*, *KAT1*, *KAT2*, and *NIP5;1.* The genes identified in this functional clade, which were mostly downregulated in MN-7WL and unchanged in SP-7WL, reveal a coordinated response underlining the plant’s ability to adapt and survive in fluctuating environmental conditions.

### Differentially expressed genes between short and long exposure to waterlogging (SP-3WL vs SP-7WL)

We next wanted to determine if pennycress plants have different responses to short (3d) versus long (7d) exposure to waterlogging. We first compared the transcriptome of SP32-10 roots waterlogged for 3 days to controls (SP-3WL) and identified 1,585 genes that were differentially expressed (554 upregulated and 1,031 downregulated). This was only about 182 fewer DEGs compared to the 7d dataset (1,767 DEGs in SP-7WL). No differential expression was detected in the leaves or silicles of SP32-10 plants waterlogged for 3 days. Next, we calculated the number of shared genes differentially expressed in the roots (compared to controls) at both 3d (SP-3WL) and 7d (SP-7WL) of waterlogging. A total of 761 genes (422 upregulated and 339 downregulated) were differentially expressed at both 3d and 7d of waterlogging, meaning more than half of the DEGs were uniquely differentially expressed at 3d (824 genes; 52%) or 7d (1,006 genes; 57%) of waterlogging (Figure 5A). Interestingly, the majority of unique DEGs present at 3d are upregulated (84%), whereas 55% of the unique DEGs at 7d are upregulated. To examine the gene function of the 761 shared DEGs between SP-3WL and SP-7WL, we conducted an over-representation analysis. Among the shared upregulated DEGs, GO terms consisted of response to hypoxia/decreased oxygen, vitamin binding, and carbohydrate kinase activity, and KEGG pathways consisted of fructose and mannose metabolism, glycolysis, MAPK signaling, and starch and sucrose metabolism (Figure 5B-C). Among the shared downregulated DEGs, GO terms consisted of water and fluid transport, lysosome, and water channel activity, and KEGG pathways consisted of ascorbate/aldarate metabolism and cyanoamino acid metabolism (Figure 5B-C). This indicates that pennycress roots under short and long-term waterlogging undergo similar stress responses, such as those involved in hypoxia and energy production.

**Figure 5.**
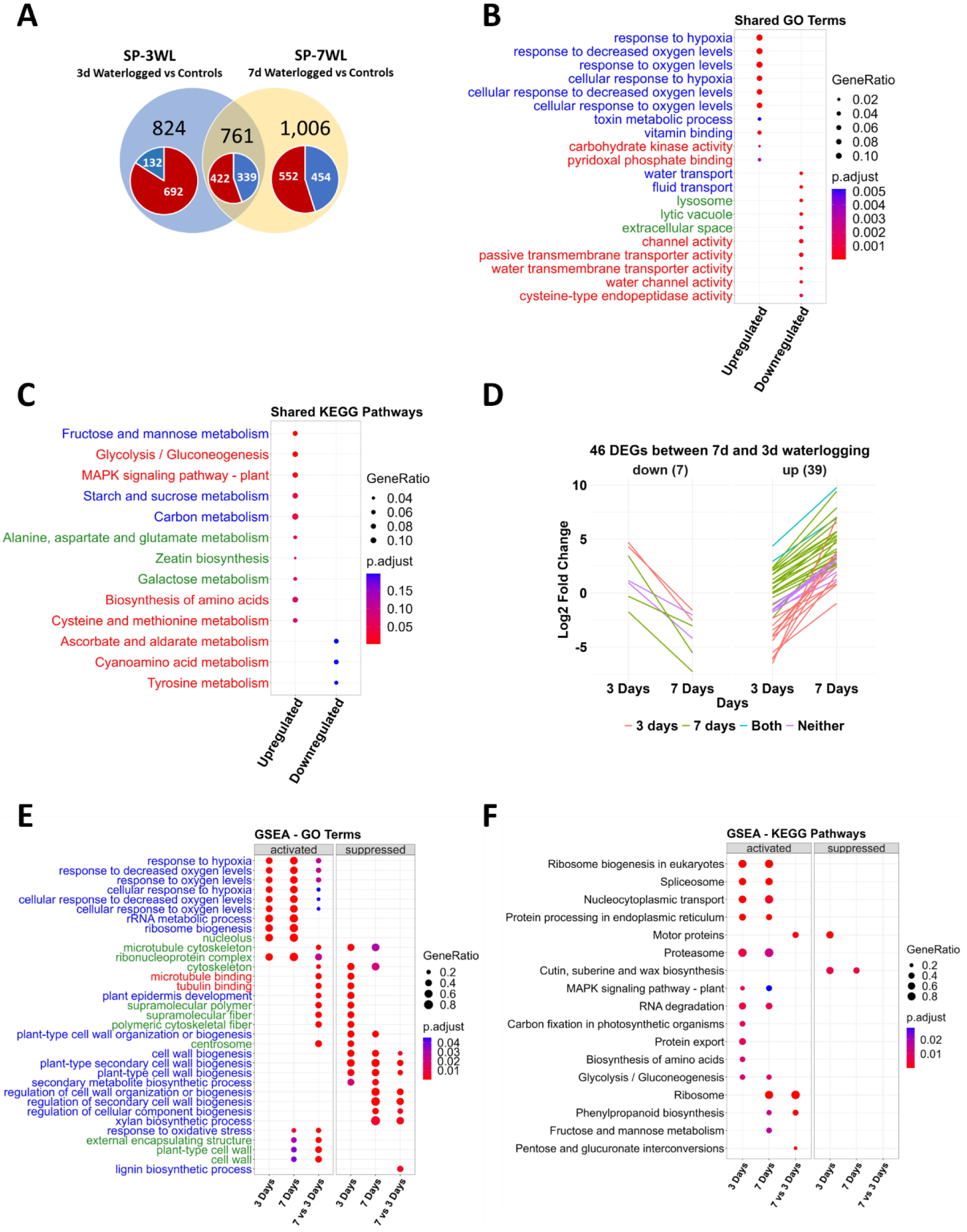
Differentially expressed genes (DEGs) and enrichment results between 7 days and 3 days of waterlogging in SP32-10 roots. (**A**) Venn diagram of overlapping DEGs from SP-3WL and SP-7WL. Red, and blue colors in pie chart represent genes up and downregulated respectively. (**B-C**) Over-representation analysis of the 10-20 most significant GO terms and KEGG pathways of the 761 shared DEGs between SP-3WL and SP-7WL. Blue, red, and green terms represent biological process, cellular component, and molecular function, respectively. GeneRatio represents the ratio of the number of genes from the input data that belong to each term to the total number of genes in the database associated with that term and p.adjust represents the level of significance of the enrichment. (D) The log2FC of the 46 DEGs from the interaction of SP-7WL vs SP-3WL. Blue-colored lines indicate genes that were significant in both SP-7WL and SP-3WL, coral and green lines indicate genes significant in only SP-3WL and SP-7WL respectively, and purple lines indicate genes that were not significant in either. (**E-F**) Gene set enrichment analysis of the 15-30 most significant GO terms and KEGG pathways of SP-3WL, SP-7WL, and the interaction of SP-7WL vs SP-3WL.

To examine genes significantly different in the roots between the time points, we tested the interaction between timepoint and treatment (SP-7WL vs SP-3WL), revealing only 46 DEGs (39 upregulated and 7 downregulated) (Figure 5D). Among these 46 DEGs, 13 were significantly differentially expressed in SP-3WL, 23 were significantly differentially expressed in SP-7WL, and 2 were significantly differentially expressed in both SP-7WL and SP-3WL (Figure 5D and Supplemental File 6). These 13 DEGs specific to SP-3WL were primarily downregulated (11), whereas the 23 DEGs specific to SP-7WL were primarily upregulated (20). In conclusion, few genes had a significant change of foldchange between 3 and 7 days of waterlogging in SP32-10, but a handful of genes did increase expression as waterlogging persisted.

Next, we performed a GSEA with the genes differentially expressed in SP-3WL, SP-7WL, and SP-7WL vs SP-3WL (Figure 5E-F and Supplemental Files 2 and 4). At 3d of waterlogging, DEGs were enriched for 354 GO terms and 21 KEGG pathways, whereas at 7d of waterlogging, DEGs were enriched in only 135 GO terms and 14 KEGG pathways. At 7d of waterlogging compared to 3d of waterlogging, DEGs were enriched for 153 GO terms and 4 KEGG pathways. Upregulated genes in SP-7WL vs SP-3WL were enriched in categories such as response to hypoxia, response to oxidative stress, and plant-type cell wall loosening. Some of the 39 upregulated interaction DEGs fell into the category of oxidative stress, such as *STACHYOSE SYNTHASE* (Δlog2FC = 5.7) and a gene encoding a heat shock protein (Δlog2FC = 4.8) (Supplemental File 6). Other DEGs were involved in cell wall modification, pectinesterase activity, carbohydrate catabolic process, and hydrolase activity, such as a gene encoding a pectin lyase-like superfamily protein (Δlog2FC = 7.5) and *PECTIN METHYLESTERASE INHIBITOR-PECTIN METHYLESTERASE 20* (Δlog2FC = 3.8). There were also a handful of GO terms related to microtubule cytoskeleton, tubulin binding, plant epidermis development, and supramolecular fiber that were more upregulated in SP-7WL than SP-3WL, because these categories were all strongly downregulated at 3d of waterlogging compared to controls, but not at 7d. Downregulated genes in SP-7WL vs SP-3WL were enriched for GO terms that were also enriched in SP-7WL downregulated genes, such as regulation of secondary cell wall biogenesis and xylan biosynthesis. Additionally, some of the 7 downregulated interaction DEGs were involved in stress responses like response to cold, water deprivation, and phosphate starvation, such as *LATE EMBRYOGENESIS ABUNDANT 4-5* (*LEA4-5*, Δlog2FC = −9) and *MONOGALACTOSYL DIACYLGLYCEROL SYNTHASE 3* (*MGD3*, Δlog2FC = −3.2). These results revealed that some stress responses, such as the response to hypoxia, were more upregulated at 7d of waterlogging compared to 3d, whereas some responses, such as cell wall biogenesis, were more downregulated (Supplemental Figure 9). Additionally, there were some genes belonging to GO terms that did not significantly change in expression between short and long periods of waterlogging, such as rRNA metabolic process, ribosome biogenesis, and the cytoskeleton. Lastly, there were very few differences in KEGG pathways enriched in SP-7WL vs SP-3WL. In conclusion, there were similar responses between long and short-term waterlogging, as genes involved in response to hypoxia were upregulated and genes involved in cell well biogenesis were downregulated at both 3d and 7d, but regulation was stronger at 7d. Unique downregulation of cytoskeleton genes was seen at 3d and not 7d.

### Differentially expressed genes following 3 hours of recovery from waterlogging

The sudden removal of a stressful condition can often trigger a stress response in plants, such as the sudden reoxygenation of roots once flood waters recede. To capture the transcriptome response of this environmental switch, gene expression was examined during the early hours (3h) of recovery from 7d of waterlogging. We investigated the DEGs between 3h recovery and controls (MN-REC and SP-REC) and found 2,398 genes were differentially expressed in MN-REC roots (1,153 upregulated and 1,245 downregulated) and 1,473 genes were differentially expressed in SP-REC roots (791 upregulated and 682 downregulated), but very few DEGs were detected in leaves and silicles during recovery (Supplemental File 1). This is 1,026 DEGs fewer than in MN-WL roots and 294 fewer than SP-WL roots. Furthermore, 2,141 root DEGs in MN106 and 1,168 root DEGs in SP32-10 were shared between 7d of waterlogging and 3h of recovery. We performed an over-representation analysis with these shared DEGs and found shared upregulated genes (1,038 genes in MN106 and 647 genes in SP32-10) were enriched for GO terms such as response to hypoxia, glycolytic process, and response to heat, as well as KEGG pathways such as glycolysis, fructose and mannose metabolism, and biosynthesis of amino acids (Supplemental Files 2 and 4). Shared downregulated genes (1,103 in MN106 and 521 in SP32-10) were enriched for GO terms such as response to extracellular stimulus, water transport, cell wall biogenesis, and glucosinolate metabolic process, as well as KEGG pathways such as plant secondary metabolite biosynthesis, plant hormone signal transduction, and terpenoid-quinone biosynthesis. This revealed that many genes involved in waterlog stress responses were still activated after the plants had been removed from the water. Comparing the recovery tissues to the waterlogged tissues, with the controls as the reference level, gives a better estimation of the differential gene expression in the 3 hours after the plants were removed from the water. This is because the recovery tissues were very similar to the waterlogged tissues (Supplemental Figure 10) and the roots were not fully dried out. To identify genes significantly differentially expressed between 3h recovery vs. controls and 7d of waterlogging vs. controls, we performed an interaction test. Only 4 upregulated genes in the roots were detected as differentially expressed between MN-REC and MN-7WL (Supplemental File 1). No differential expression between SP-REC and SP-7WL roots was detected.

Although very few DEGs were detected between MN-REC vs MN-7WL and SP-REC vs SP-7WL, looking at the change in expression of every gene between recovery and waterlogging through a GSEA still revealed enriched GO categories and KEGG pathways. In MN-REC vs MN-WL, genes were enriched for 94 GO categories and 19 KEGG pathways, whereas SP-REC vs SP-WL had genes enriched for 193 GO categories and 9 KEGG pathways (Supplemental Files 2 and 4). Roots after 3h of recovery compared to waterlogged roots in both accessions revealed upregulated genes enriched for GO categories such as ribonucleoprotein complex, secondary metabolite biosynthesis, and ncRNA/rRNA processing, while downregulated genes were enriched for categories such as response to decreased oxygen/hypoxia, defense response, and response to heat and chitin. Additionally, genes attributed to several KEGG pathways were upregulated in recovery roots compared to waterlogged roots, such as ribosome, biosynthesis of amino acids, carbon metabolism, and phenylpropanoid biosynthesis. On the other hand, genes related to protein processing in the endoplasmic reticulum were downregulated in recovery compared to waterlogged roots. These results indicate that in the couple of hours after the plants were removed from the waterlogged environment, some initial stress responses were lowered, such as the response to hypoxia (Supplemental Figure 11A-D), and other cellular and metabolic processes were increased, such as secondary metabolite biosynthesis (Supplemental Figure 11E-H). Therefore, while many of the same genes were differentially expressed at 3h of recovery compared to 7d of waterlogging, these genes had slight changes (less than a log2FC = |1|) in expression levels.

### N-degron pathway gene expression

Hypoxia-response genes, such as those involved in energy production under oxygen deficit, are upregulated by the ERF-VII transcription factor family which is stabilized under hypoxia via the N-degron rule pathway (Figure 6A) (Mustroph et al., 2009; Gibbs et al., 2011, 2015). This family consists of five members in Arabidopsis (*HRE1*, *HRE2*, *RAP2.12*, *RAP2.2*, and *RAP2.3*) (Giuntoli and Perata, 2018). The genes encoding HRE1 (TAV2_LOCUS15620) and HRE2 (TAV2_LOCUS13098) were significantly upregulated in both MN-7WL and SP-7WL roots by a log2FC of about 3 in MN-7WL and 4 in SP-7WL for *HRE1* and about 9 in MN-7WL and 7 in SP-7WL for *HRE2* (Figure 6A). *HRE1* and *HRE2* were also expressed at similar levels after 3d of waterlogging in SP32-10. Interestingly, the log2FC of *HRE2* was twice that of *HRE1*. *RAP2.3* (TAV2_LOCUS11345) was only differentially expressed (upregulated) in MN-7WL roots, but not in SP-7WL or SP-3WL roots. Although not significantly differentially expressed in waterlogged compared to control roots, *RAP2.12* (TAV2_LOCUS1421) was upregulated in MN106 but downregulated in SP32-10 Lastly, *RAP2.2* (TAV2_LOCUS10019) showed no significant differential expression in the waterlogged roots but was slightly upregulated in both accessions compared to controls. Furthermore, none of the *ERF-VII* genes were significantly differentially expressed between the two accessions (MN-7WL vs SP-7WL) under waterlogging.

**Figure 6.**
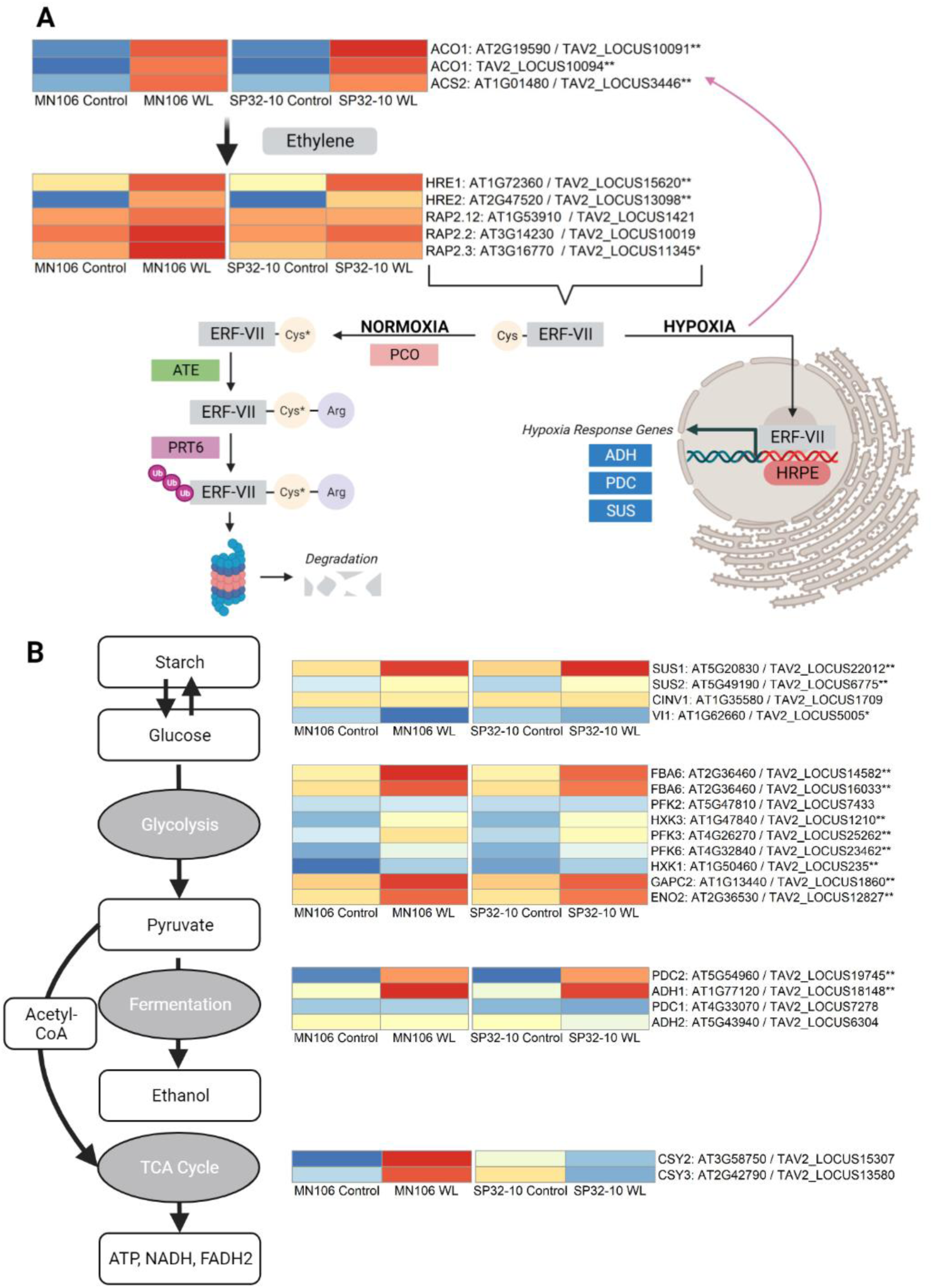
Heatmap of genes involved in the N-degron pathway and core anaerobic processes. (**A**) Genes involved in ethylene synthesis upregulate the *ERF-VII* gene family which, under normal oxygen conditions (normoxia), are susceptible to the N-degron pathway where plant cysteine oxidases (PCO) oxidate the N-terminus cysteine residue, followed by arginylation by (ATE) and tagged for ubiquitination by proteolysis 6 (PRT6) for degradation. Under hypoxia, the *ERF-VII* genes remain stable and bind to hypoxia response promoter elements of genes involved in anaerobic responses such as (**B**) starch and glucose biosynthesis/degradation, glycolysis, fermentation, and the TCA cycle. Heatmaps were made with no scale applied, meaning that scale/colors were assigned based on all values in each individual heatmap and not by row or column. A single asterisk next to the gene name indicates differential expression in waterlogged compared to control roots in MN106, and double asterisks indicate differential expression in both MN106 and SP32-10 waterlogged compared to control roots. Figure adapted from Combs-Giroir and Gschwend, 2024.

Under low oxygen, ERF-VII transcription factors bind to hypoxia response promoter elements (HRPE) of core hypoxia response genes (Licausi et al., 2010; Gasch et al., 2016), such as those involved in sugar metabolism (*SUS*), glycolysis, and fermentation (*ADH, PDC*). In MN-7WL and SP-7WL roots, *PDC2* and *ADH1* were in the top 10 most significant differentially expressed upregulated genes with a log2FC of about 8.5 and 6, respectively (Supplementary Table 5). These two genes were also among the top 10 most significant differentially expressed upregulated genes in SP-3WL roots with similar expression levels. Genes involved in the glycolysis pathway were enriched in both MN-7WL and SP-7WL roots, with upregulated genes such as *FBA1* and *FBA6*, *PFK3* and *PFK6*, and *HXK3,* with log2FC’s of about 4-5 (Figure 6B, Supplemental File 3). *SUS1-like* (TAV2_LOCUS22012) was upregulated about 4-log2FC in MN-7WL, SP-7WL, and SP-7WL roots (Figure 6B). *SUS2-like* (TAV2_LOCUS6775) was also upregulated about 3-log2FC in MN-7WL and SP-7WL roots and about 4-log2FC in SP-3WL roots. Lastly, expression levels of a cytosolic invertase (*CINV1/*AT1G35580/TAV2_LOCUS1709) were slightly downregulated in waterlogged roots in both accessions, suggesting the favoring of sucrose synthase over invertase under waterlogging (Supplemental File 1).

## Discussion

### Waterlogging caused reduced growth and impacted seed yield and oil quality

At maturity, waterlogged plants displayed a reduction in biomass and yield, as well as changes in morphology and physiology, suggesting a halt in pennycress growth as a result of the severe energy crisis invoked by waterlogging. Additionally, more morphological traits were impacted by waterlogging in SP32-10 than MN106 immediately after waterlogging, during recovery, and at the time of harvest, indicating phenotypic variation in waterlogging responses among the two accessions. Many waterlogging studies in *Brassicaceae* species have also reported reduced biomass under waterlogging, such as with *Brassica napus* (Zou et al., 2014; Wollmer et al., 2018; Liu et al., 2020; Liu and Zwiazek, 2022), *Brassica rapa* (Daugherty and Musgrave, 1994; Daugherty et al., 1994), *Brassica oleracea* (Issarakraisila et al., 2007; Casierra-Posada and Peña-Olmos, 2022), and *Arabidopsis thaliana* (Pigliucci and Kolodynska, 2002). The common reduction in plant growth and development under abiotic stress, such as flooding, could be attributed to multiple factors, such as nutrient deficiency (Steffens et al., 2005), the trade-off with mounting a defense response (Bechtold and Field, 2018), and reduced ATP synthesis and photosynthetic rates (Bailey-Serres and Voesenek, 2008; Xu et al., 2019). While biomass was not significantly reduced in either accession directly after waterlogging, at the time of harvest, root dry weight was significantly reduced in SP32-10, but remained unchanged in MN106. It’s possible that MN106 utilized a quiescence strategy under waterlogging by halting growth to conserve energy until the water receded, similar to *Rorippa sylvestris*, lowland rice cultivars, and *Rumex acetosa* under full submergence (Fukao et al., 2006; Akman et al., 2012; van Veen et al., 2013), and with some evidence in *B. napus* and *A. thaliana*, as well (Lee et al., 2011; Wittig et al., 2021). This is supported by the observation of many downregulated genes related to cell growth and the cell wall in MN-7WL compared to SP-7WL roots. Oppositely, after waterlogging SP32-10 had significantly reduced root and shoot biomass, more dead inflorescences after 1 and 2 weeks of recovery, and matured more rapidly, indicating that this accession might be less equipped to survive waterlogging stress. This is supported by the fact that SP32-10 had reduced seed yield in both waterlogging experiments unlike MN106.

Additionally, our transcriptome data revealed several differences between the accessions under waterlogging that might contribute to differences in biomass and yield. Genes within many GO categories and pathways related to cell wall biogenesis, plant epidermis development, and the cell cycle were significantly downregulated in waterlogged plants, indicating growth was halted. Waterlogging at the germination stage in *Brassica napus* also showed downregulation of “starch and sucrose metabolism” and “photosynthesis” related genes that can explain the observed reduced development of the hypocotyl, shoot, and roots, as well as differential expression of several genes that might play a role in cell differentiation and division in the root apical meristem, such as *IAA17* and *ACR4* (Guo et al., 2020). In MN-7WL roots, genes enriched for root epidermal cell differentiation were downregulated and an *IAA17*-like gene (*TAV2_LOCUS6916*) and *ACR4*-like gene (*TAV2_LOCUS15261*) were significantly differentially expressed (down and upregulated respectively), suggesting the inhibition of root development. This was not observed in SP32-10 waterlogged roots. Other downregulated genes involved in root growth and development were a *MIZ* gene, *BAM3*, and *SKS15*. The *MIZ1* gene in Arabidopsis, a representative of the MIZU-KUSSEI-like (MIZ) proteins, is essential for root hydrotropism, or the directional growth of roots toward moisture gradients. *MIZ1*encodes a protein of unknown function, but it is expressed in the lateral root cap and cortex of the root proper and is localized as a soluble protein in the cytoplasm and in association with the endoplasmic reticulum (ER) membranes in root cells. The expression of *MIZ1* is regulated by abscisic acid (ABA), influencing the hydrotropic response. Overexpression of *MIZ1* resulted in an enhancement of hydrotropism, thus, in a waterlogged state the suppression of hydrotropism via downregulation of *MIZ1* is a logical response (Moriwaki et al., 2011). Similarly, *BAM3* is essential in protophloem differentiation and root meristem growth. It acts as a receptor-like kinase that interacts with specific peptide ligands. Mutations in *BAM3* can suppress the root meristem growth and perturb protophloem development, indicating its critical role in these processes (Depuydt et al., 2013), so downregulation of *BAM3* is consistent with a halting of hydrotropism. In addition, *SKU5*, a homolog of *SKS15*, is known to play a role in affecting directional growth processes (Sedbrook et al., 2002). Recent evidence suggests that *SKU5* is involved in reactive oxygen species (ROS) homeostasis and that planes of cell division, thus directional growth cues, are intrinsically related to levels of ROS within the root cell apoplastic environment (Chen et al., 2023). The increased expression of *MATE1* we observed in this study is consistent with increased accumulation of ROS within root cells. Loss of function mutants (*sku5*) have shown increased ROS signaling, aberrant planes of cell division, and display a characteristic root growth patterns that shifts away from the gravity vector (Sedbrook et al., 2002). If this pennycress *SKS15* gene functions similarly to *SKU5*, it can be reasoned that reduced expression of this gene could trigger the ROS signaling cascade that allows lateral roots to explore soil layers closer to the surface that are less anoxic during waterlogging granting a short-term adaptive advantage.

A primary concern of flooding events is the impact on seed yield, oil content, and oil quality. Waterlogging studies on *B. napus* at the stem elongation and early flowering stages have reported yield loss and/or negative impacts on oil quality (Xu et al., 2015; Wollmer et al., 2018), and sensitivity to waterlogging at the flowering stage has also been reported in other crops such as soybean (Wu et al., 2017) and sesame (Wang et al., 2021). Seed oil content was not significantly reduced after waterlogging in the greenhouse experiment, indicating that oil yield was not affected by waterlogging. In MN106 waterlogged seeds from the greenhouse experiment, there were several changes in some of the major fatty acid constituents found in pennycress seed oil (palmitic, linoleic, erucic, and eicosenoic acid), indicating different oil properties could be impacted by waterlogging, but the consequence of this depends on the end-use of the oil. However, there were very few changes in the oil profiles of MN106 waterlogged seeds from the growth chamber experiment. These discrepancies between experiments could be due to environmental conditions and/or harvest dates since greenhouse temperatures were about 2° warmer and plants were harvested about 2-3 weeks later in the greenhouse. In the RNA-seq dataset, upregulated DEGs in MN-7WL silicles were enriched in functions related to response to fatty acid, cellular response to lipid, fatty acid catabolic process, lipid modification, and lipid oxidation, and downregulated DEGs enriched for fatty acid biosynthesis. Oppositely, SP-7WL silicle DEGs did not have GO category enrichments. For MN-7WL silicles, genes involved in the synthesis of very long-chain fatty acids (VLCFAs) were some of the core enriched genes in the fatty acid biosynthesis and fatty acid catabolic category, such as genes encoding 3-ketoacyl-CoA synthase enzymes, plant natriuretic peptide (PNP), ELO homolog 2, and Acyl-CoA oxidase 1 and 2. This could be the reason for decreased levels of very long-chain fatty acids in the seed oil of MN106 waterlogged plants, such as eicosenoic, behenic, and erucic acid, which was not observed in SP32-10.

### Waterlogging caused changes in energy production and activation of core hypoxia response genes

Under flooding and/or hypoxia stress in plants, mitochondrial respiration is severely compromised by a lack of oxygen, causing an increase in glycolysis, ethanol fermentation, and sugar metabolism (Sairam et al., 2008). The accumulation of pyruvate from glycolysis is converted to acetaldehyde by PDC, and ADH then reduces acetaldehyde to ethanol to facilitate the regeneration of NAD+ from NADH. While this helps to maintain energy production, it is an inefficient method because only 2 ATP are generated compared to approximately 36 ATP produced under aerobic respiration. In the roots of both accessions under waterlogging, *PDC2* and *ADH1* were very highly differentially expressed, and genes involved in glycolysis were upregulated. Additionally, upregulated genes involved in the generation of precursor metabolites and energy were significantly enriched in MN106 compared to SP32-10 waterlogged roots, which were mostly glycolysis genes such as *FBA6*, *FBA3*, *GAPC2*, and *PCK2.* This showcases the ability of MN106 to strongly upregulate glycolysis in waterlogged roots more than SP32-10 to sustain energy production.

Aside from glycolysis, we detected many differences in gene expression between MN106 and SP32-10 under waterlogging. Compared to the MN106 draft genome, SP32-10 had about 410,000 variants, with about 20,000 non-synonymous variants residing in about 23% of the genes (McGinn et al., 2019). This could explain some of the phenotypic and genetic variation we observed between the accessions. For instance, genes related to hypoxia/decreased oxygen were significantly upregulated in MN106 waterlogged roots compared to SP32-10 waterlogged roots (Figure 4C-E). Some of the genes contributing to the core enrichment of these categories were *ATPS2, SAG14*, and *RBOHD*. *SAG14* is known to play a role in leaf senescence (Hao et al., 2022; Hsu et al., 2022) and is induced under limited phosphate conditions (Linn et al., 2017). In *A. thaliana*, RBOHD, an NADPH oxidase, plays a role in the response to oxygen deficiency (Pucciariello et al., 2012; Gonzali et al., 2015; Yang and Hong, 2015; Liu et al., 2017), as well as waterlogging (Sun et al., 2018) and submergence (Hong et al., 2020), through production of reactive oxygen species, such as H_2_O_2_, and increased activity of hypoxia-induced genes like *ADH* and *PDC*. Both *RBOHD* and *SAG14* are activated by the WRKY transcription factor gene *WSR1* in *B. napus*, which contributes to cell death and leaf senescence (Cui et al., 2020). Furthermore, some genes encoding proteins of the pyruvate kinase/phosphate dikinase substrate cycle, which synthesizes inorganic pyrophosphate (Ppi), were significantly upregulated in waterlogged roots of both accessions, but slightly more upregulated in MN106 than SP32-10. This could indicate Ppi production for glycolysis and a mechanism for improved waterlogging tolerance. For instance, anoxia-tolerant species, such as rice, have been shown to use Ppi as high-energy donor molecules under ATP deficiency, where Ppi can be synthesized from the pyruvate kinase/phosphate dikinase substrate cycle or in the reverse reaction of membrane-bound H^+^-pyrophosphatase (Huang et al., 2008; Igamberdiev and Kleczkowski, 2021). Oppositely, the absence of Ppi-dependent glycolysis has been observed in *Arabidopsis thaliana*, an anoxia-intolerant species (Loreti et al., 2005; Igamberdiev and Kleczkowski, 2021). Interestingly, gene clusters differentially regulated between two flooding-tolerant *Rorippa* species and *A. thaliana* exposed genes associated with this pathway, indicating this could be a shared flooding tolerance mechanism to support energy metabolism within the *Brassicaceae* family (Sasidharan et al., 2013). In MN106 waterlogged roots, an inorganic pyrophosphatase, *ATPS2*, was among the top 10 significant differentially expressed genes and was one of the top three significantly differentially expressed genes between MN106 and SP32-10 waterlogged roots. In Arabidopsis, *ATPS2* is highly expressed under phosphate starvation (Baek et al., 2013). Consequently, *ATPS2*, and (Ppi)-dependent alternative pathways of phosphorylation, are worth future exploration for improved waterlogging tolerance in pennycress.

Another expected hypoxic response was the differential expression of *SUS1* and *SUS2* in the roots of both pennycress accessions under waterlogging. Elevated sucrose synthase transcript and protein abundance were also detected in response to flooding and hypoxia in several species such as wheat (*Triticum aestivum*) (Albrecht and Mustroph, 2003), potato (*Solanum tuberosum*) (Biemelt et al., 1999), *Arabidopsis thaliana* (Loreti et al., 2005; Bieniawska et al., 2007), rice (*Oryza sativa*) (Locke et al., 2018), cucumber (*Cucumis sativus*) (Kęska et al., 2021), and zombi pea (*Vigna vexillate*) (Butsayawarapat et al., 2019). Sucrose synthase, which catalyzes the reversible cleavage of sucrose into fructose and UDP or ADP-glucose, is suggested to play a beneficial role in alleviating anaerobic stress by providing an energetically efficient method of supplying UDP-glucose for biological processes, whereas sucrose hydrolysis via invertases requires several steps to produce UDP-glucose (Stein and Granot, 2019). This has been supported by the observation of reduced invertase activity under low oxygen (Albrecht and Mustroph, 2003; Bieniawska et al., 2007). Sucrose synthase mutants have also been observed to perform poorly under flooding or hypoxic growth conditions (Bieniawska et al., 2007; Yao et al., 2020), further supporting its involvement under oxygen deficiency.

Waterlogged roots of both pennycress accessions displayed a strong induction of genes related to the ribosome, ribonucleoprotein, and translation compared to controls. Interestingly, these processes were downregulated in the leaves. Since protein synthesis is an energetically costly process, it is surprising to see genes involved in these processes enriched when ATP production is so low under hypoxia, but this hints at the ability of pennycress to initiate stress-specific responses, such as activating genes encoding proteins involved in glycolysis, anaerobic fermentation, and signaling pathways. For instance, selective mRNA translation under hypoxia could be a method for energy conservation under low oxygen stress (Branco-Price et al., 2008).

### Waterlogging caused upregulation of ERF-VII transcription factors

Hypoxia-response genes, such as those involved in energy production under oxygen deficit, are upregulated by the ERF-VII transcription factor family. The genes encoding ERF-VII members HRE1 and HRE2 were significantly upregulated in waterlogged roots in both accessions, with *HRE2* doubled in expression compared to *HRE1* and more upregulated in MN106 than SP32-10 (1.7 log2FC difference). Previous research has shown that *HRE1* requires protein synthesis of transcription factors to be induced, whereas *HRE2* is likely to be constitutively expressed, but its mRNA is unstable under normoxia (Licausi et al., 2010). The same study also proposed that *HRE1* might upregulate *HRE2*. This could be a potential reason for why *HRE2* is more strongly upregulated under waterlogging in pennycress. Strong induction of *HRE2* was also found in dark-submerged *Rorippa* roots (van Veen et al., 2014) and hypoxic Arabidopsis roots (Licausi et al., 2010), possibly indicating a conserved response across *Brassicaceae* when roots are exposed to hypoxic environments. Additionally, *RAP2.3* was significantly upregulated in waterlogged MN106 roots, but not in SP32-10. This higher expression of *ERF-VIIs* (*HRE2* and *RAP2.3*) in MN106 compared to SP32-10 suggests greater transcriptional control over downstream hypoxia-response genes. Like most other plant species, pennycress exhibits expression of *ERF-VII* genes under waterlogging, in turn upregulating common hypoxic responses such as glycolysis and fermentation.

### Waterlogging resulted in differential expression of genes involved in root cell wall components between MN106 and SP32-10

Under waterlogging, plants may be exposed to increased iron availability in the soil, leading to the absorption of excess iron (Lapaz et al., 2020). Therefore, the strong downregulation of *bHLH100*, a key regulator of iron-deficiency responses (Sivitz et al., 2012), and *IRT1*, a member of the ZIP metal transporter family (Henriques et al., 2002), in MN106 roots might be a response to iron excess or toxicity. Iron toxicity in plants under flooding has not been well-studied, but under iron excess, there are mechanisms to prevent uptake and oxidative stress within the cell, such as casparian strips (Barberon et al., 2016; Oliveira de Araujo et al., 2020; Lapaz et al., 2022). Casparian strips are apoplastic barriers composed of lignin in a ring-like structure around endodermal cells (Naseer et al., 2012). Interestingly, other significant downregulated DEGs in MN106 roots compared to SP32-10 were peroxidases, which are required for casparian strip lignification (Rojas-Murcia et al., 2020) and a putative casparian strip membrane protein. Furthermore, genes downregulated in MN106 compared to SP32-10 were enriched for casparian strip and secondary cell wall categories. Defects in the casparian strip, caused by loss of function mutations in *CASP1*/*CASP3*, *ESB1*, *GASSHO1*, and *MYB36*, led to compensatory lignification, where the middle lamellae of cell corners became lignified and casparian strips were unfused (Roppolo et al., 2011; Hosmani et al., 2013; Kamiya et al., 2015; Doblas et al., 2017; Nakayama et al., 2017; Barbosa et al., 2019). The downregulation of these casparian strip genes indicates that waterlogged MN106 roots might have had disrupted, or discontinuous, casparian strips and reduced lignin deposits in the root cells which could affect nutrient homeostasis or apoplastic transport. This might help explain the strong downregulation of genes related to iron levels. It is also possible that the downregulation of genes involved in casparian strip formation and lignification is a result of reduced lignin content in the roots. Reductions in lignin content have been observed in plants under waterlogging or submergence such as wheat (Nguyen et al., 2016) and soybean (Komatsu et al., 2010). Soluble sugars, specifically glucose, are in demand under waterlogging, which potentially sends more phosphoenolpyruvate to glycolysis rather than the shikimate pathway where the precursor for lignin, phenylalanine, is produced (Vanholme et al., 2012). Therefore, the downregulation of lignin genes in the waterlogged roots might instead be a response to decreased oxygen and low ATP production in the roots, increasing the need for phosphoenolpyruvate to support glycolysis.

The downregulation of genes involved in lignin and suberin biosynthesis in waterlogged MN106 roots was an unexpected result since these components can serve beneficial roles under flooding, such as formation of radial oxygen-loss (ROL) barriers to prevent oxygen leakage and entry of soil-derived gases and phytotoxic substances (Ejiri et al., 2021; Jiménez et al., 2021). However, the downregulation of these genes might be a mechanism for conserving growth and energy in response to hypoxia. Suberin is composed of fatty acids, such as VLCFAs, and it has been shown that rice roots forming ROL barriers under stagnant conditions accumulate malic acid and VLCFAs (Kulichikhin et al., 2014). Therefore, one explanation for reduced suberin biosynthesis in MN106 waterlogged roots is the preferential breakdown of VLCFAs for ATP production, which is supported by the fact that fatty acid beta-oxidation was up-enriched in MN106 waterlogged roots compared to SP32-10. Furthermore, HXXXD-type acyl-transferase family proteins are involved in the biosynthesis of lipid polymers that are components of the cuticle and suberized cell walls. These structures are crucial for preventing dehydration and providing a physical barrier against pathogens. A gene encoding this protein was downregulated in MN106 roots during waterlogging, which could be part of a broader metabolic adjustment to prioritize energy conservation and reallocation of resources to more critical survival processes, such as anaerobic respiration (Molina and Kosma, 2015). Also involved in lipid metabolism, GGL25 is part of the larger family of GDSL-type esterase/lipase genes in plants (Lai et al., 2017). Genes encoding GGL25 and a GDSL-motif esterase/acyltransferase/lipase were more downregulated in MN106 than SP32-10 roots, indicating a role in osmotic stress resistance (Ding et al., 2019) and involvement in the biosynthesis of suberin (Klein, 2019). Another study involving AT4G24160, a close homolog to *GGL25* in Arabidopsis, revealed that it is an acyl-CoA-dependent lysophosphatidic acid acyltransferase with additional enzymatic activities, including triacylglycerol lipase and phosphatidylcholine hydrolyzing activities (Ghosh et al., 2009). This suggests that GGL25 might play a pivotal role in maintaining lipid homeostasis in plants by regulating both phospholipid and neutral lipid levels. In addition, AT4G24140, an alpha/beta hydrolase enzyme, may be involved in the biosynthesis of fatty acids and suberin (Compagnon et al., 2009). Considering their functional roles and connection to suberin, it is plausible that GDSL-type esterase/lipases, HXXXD-type acyl transferases, and an alpha/beta hydrolase (AT4G24140) could be involved in related or complementary pathways in lipid metabolism and modification of extracellular structures such as suberin in plants (Figure 7).

**Figure 7.**
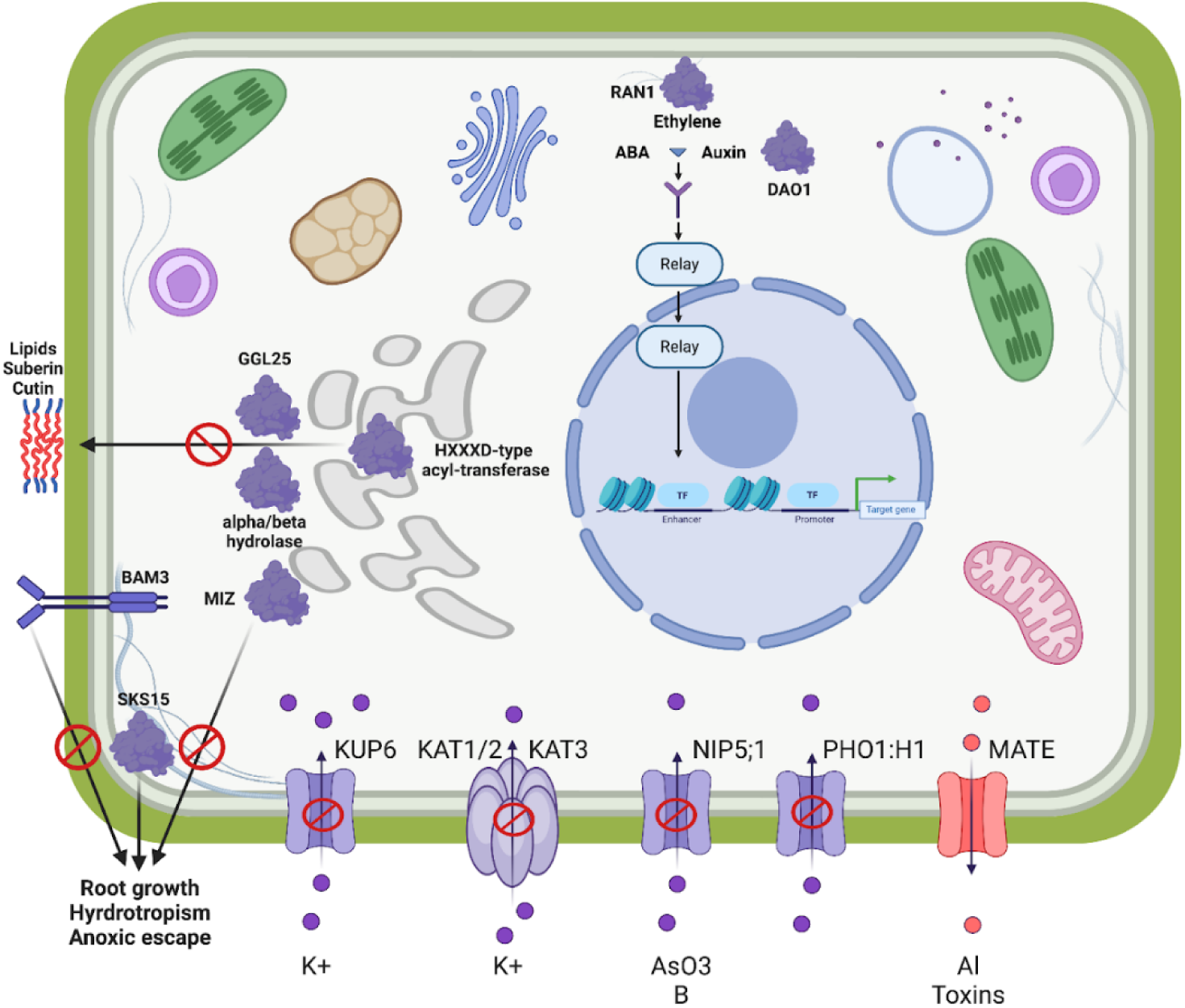
Conceptual model of a differentially expressed functional clade of genes. The downregulation of inward transport associated with KUP6, KAT3, NIP5:1, and PHO1;H1 and upregulation of eflux pump MATE helps to maintain osmotic balance and cell health. The downregulation of a HXXXD-type acyl transferase, alpha/beta hydrolase and GGL24 will decrease the production of suberin/cutin since water conservation is not required during waterlogging. Similarly, MIZ, SKS15, and BAM3 are downregulated as hydrotropism is not required during waterlogging. Hormones play an active role in the transcriptional regulation of many of these genes, including RAN1, which regulates the ethylene response pathways and DAO1 that affects auxin levels.

The differential expression of genes related to the primary cell wall was also observed between accessions under waterlogging. For instance, *XYLOGLUCAN ENDOTRANSGLUCOSYLASE*, *PECTATE LYASE,* and *PECTINESTERASE* genes, which are involved in cell wall degradation and loosening, were significantly upregulated in MN106 waterlogged roots compared to SP32-10. However, a gene with a similar function, *EXPANSIN 18*, was significantly downregulated in MN106. These cell wall-related proteins are involved in the modification of cell wall extensibility, which facilitates cell enlargement and elongation (Sampedro and Cosgrove, 2005; Eklöf and Brumer, 2010; Sénéchal et al., 2014), and have been implicated in flooding stress in other crops such as soybean (Nanjo et al., 2011, 2013), wheat (Kong et al., 2010), maize (Vitorino et al., 2001), and cucumber (Kęska et al., 2021). This was mostly observed as a downregulated response likely corresponding with root growth inhibition under waterlogging. Interestingly, similar to our expression results, XTH proteins were decreased in abundance in a sensitive rapeseed variety under waterlogging but increased in a tolerant variety, with root growth suppressed in both, but more so in the sensitive variety (Xu et al., 2018). Additionally, overexpression of an *XTH* gene in soybean enhanced waterlogging tolerance and increased root growth (Song et al., 2018). Unlike SP32-10, we observed reduced root weight in MN106 immediately after waterlogging, indicating that these genes might not be upregulated for root cell elongation, or that root growth did not precisely coincide with these gene expression changes. The fact that MN106 waterlogged plants did not have significant changes in root biomass at maturity supports the idea that the phenotype presented itself later. Transcriptional alterations in these cell wall genes might be an adaptive mechanism in MN106 to better withstand waterlogging through other traits like aerenchyma formation. For instance, cell wall modification plays an important role in aerenchyma formation in maize and wheat (Arora et al., 2017; Li et al., 2019).

### Genes involved in water and ion homeostasis were downregulated in waterlogged MN106 roots

Another unique role between MN106 and SP32-10 under waterlogging was the downregulation of genes involved in water and ion homeostasis in MN106. It is likely that an important component of the adaptive response to waterlogging is the regulation of potassium homeostasis, a key element in maintaining cell turgor and osmotic balance (Osakabe et al., 2014). The downregulation of the *KUP6* potassium transporter in MN106 is thus likely to be a significant player and is directly regulated by abscisic acid (ABA) signaling pathways. Complementing this is the downregulation of the *KAT3* gene, also known as *AtKC1*, which is part of the Shaker family of voltage-gated potassium channels in *Arabidopsis thaliana*. It has been shown that KAT3 does not form functional potassium channels on its own. Instead, it serves as a modulator of other Shaker family channel activities, such as those mediated by KAT1, KAT2, and AKT2. Specifically, KAT3 can form heteromeric channels with these inward-rectifying channels, altering their conductance and shifting their activation potential (Jeanguenin et al., 2011). KAT1, an inward-rectifying channel, is pivotal for the inward transport of potassium ions (K+) into the cells, resulting in water influx through osmosis, causing cells to swell (Schachtman et al., 1992; Uozumi et al., 1995; Saponaro et al., 2017; Locascio et al., 2019). They play a crucial role in maintaining potassium homeostasis. Thus, the downregulation of *KAT3* in MN106 would be a logical response to waterlogging stress.

In MN106 waterlogged roots, the downregulation of *NIP5;1*, a member of the Major Intrinsic Protein (MIP) family, including Nodulin-26-like Intrinsic Proteins (NIPs), may also play a role in this story. These proteins, acting as aquaporins, are crucial for regulating water and ion transport across cellular membranes. The downregulation of *NIP5;1* during waterlogging stress could be a mechanism to maintain osmotic balance by controlling levels of boron, as NIP5;1 facilitates the transport of boric acid in addition to water. Under conditions of boron deficiency, NIP5;1 is crucial for boron uptake, playing a significant role in plant growth and development (Takano et al., 2006). The interaction between salinity and boron affects root hydraulic conductance and nutrient uptake, and is mediated by changes in aquaporin expression, including *NIP5;1* (Martinez-Ballesta et al., 2008). Moreover, NIP5;1 is permeable to arsenite, and changes in its expression could influence arsenite transport (Mitani-Ueno et al., 2011). The downregulation of *NIP5;1* during waterlogging might therefore be a regulatory mechanism to modulate the internal levels of boron and arsenite, maintaining osmotic and ionic balance under stress conditions.

The *PHO1;H1* gene plays a crucial role in inorganic phosphate (Pi) transport and homeostasis. PHO1;H1 contributes to Pi loading into the xylem and is regulated by Pi deficiency via distinct pathways (Stefanovic et al., 2007). The downregulation of *PHO1;H1* in MN106 may be linked to the plant’s osmotic stress response, adjusting its physiological processes to cope with the excess water in the soil environment. Both auxin and ABA are known to regulate *PHO1;H1* expression (Ribot et al., 2008). Collectively, *KUP6*, *KAT3*, *PHO1;H1*, and *NIP5;1* likely contribute to ion balance and osmotic stability, with their downregulation suggesting a critical role in the plant’s adaptation to waterlogged soils (Philippar et al., 2004). Furthermore, *DAO1*, involved in auxin metabolism, plays a nuanced role in this scenario. The regulation of *DAO1* expression is vital for maintaining auxin at optimal levels, crucial for plant growth and development, particularly in stress conditions like waterlogging (Mellor et al., 2016).

A gene encoding a MATE Efflux Family Protein was the only gene upregulated in MN106 in the functional clade examined from the MENTOR results. MATE proteins are known for their ability to transport a wide range of substrates, including organic compounds, plant hormones, and secondary metabolites, out of cells. This transport mechanism plays a critical role in various plant processes and stress responses. In the context of waterlogging stress, the upregulation of MATE efflux proteins can be attributed to several factors, including 1) Detoxification and Stress Response: MATE proteins are involved in the detoxification of toxic compounds and xenobiotics. During waterlogging, plants may experience an accumulation of toxic metabolites due to reduced oxygen availability and altered metabolic processes. The upregulation of a MATE under such conditions could be a protective mechanism to remove harmful compounds from the cells, thereby aiding in stress tolerance and survival (Liu et al., 2009). 2) Hormonal Regulation and Homeostasis: MATE proteins are also involved in transport of plant hormones. Waterlogging stress alters the hormonal balance in plants, impacting growth and development. The upregulation of MATE efflux proteins may be part of the mechanism to maintain hormonal homeostasis under stress conditions (Upadhyay et al., 2020). 3) Response to Abiotic Stresses: Studies have shown that MATE genes are responsive to various abiotic stresses, including aluminum toxicity, which can be exacerbated under waterlogged conditions. The upregulation of this gene might be a response to the increased aluminum availability in acidic soils during waterlogging, helping plants to tolerate such adverse conditions (Duan et al., 2013). The upregulation of MATE efflux family proteins during waterlogging stress is likely an adaptive response to manage the internal cellular environment and help the plant mitigate the adverse effects of waterlogging to maintain its physiological functions (Figure 7).

### The potential role of hormones under waterlogging stress in pennycress

Abscisic acid (ABA), auxin, and ethylene play pivotal roles in facilitating survival and growth in waterlogged plants. ABA signaling pathways are important in maintaining cellular integrity during waterlogging; They regulate KUP6 (Osakabe et al., 2014) and PHO1;H1 (Ribot et al., 2008), as discussed above, which are important in balancing potassium and phosphate levels and osmotic adjustment under osmotic stress conditions common in waterlogging scenarios. Furthermore, auxin regulates a variety of morphological and anatomical responses to flooding stress. It acts synergistically with ethylene to control adventitious root and aerenchyma formation, enhancing root plasticity and adapting the root architecture to waterlogged conditions for better survival (Hu et al., 2016). The interaction between auxin and other hormones like ABA and GA is crucial in controlling morphological responses such as shoot elongation, root formation, and stomatal opening and closure under waterlogged conditions (Hu et al., 2016). *KAT3*-associated proteins, potassium channels *KAT1* and *KAT2,* are regulated by auxin (Ribot et al., 2008). The expression of *MIZ1* in roots, critical for hydrotropic response, may be influenced by auxin levels, indicating a possible modulation of root growth direction in response to auxin under waterlogged conditions (Moriwaki et al., 2011). Ethylene is known to play a central role in most morphological and anatomical responses of plants to flooding. It is involved in controlling aerenchyma formation, adventitious root formation, and shoot elongation under flooded conditions. The interaction between ethylene and other phytohormones such as auxin, ABA, and GA is critical in regulating these responses (Hu et al., 2016). *RAN1* was downregulated in MN106 under waterlogging and it is known that loss-of-function mutation in *RAN1* can result in constitutive activation of the ethylene response pathway (Woeste and Kieber, 2000). We note that Nodulin-26-like Intrinsic Proteins (NIPs) expression can be influenced by ethylene. The increase in ethylene under such conditions can influence the expression of various genes involved in water and oxygen stress responses, as discussed in the above section (Loreti et al., 2016). The interactions of these hormones are complex and crucial in regulating plant growth and stress responses. Each of these hormones has distinct roles, but their interplay is what allows plants to adapt effectively to the challenges of excess water and reduced oxygen availability.

### Future perspectives

In conclusion, waterlogging in pennycress during the reproductive growth phase can impair growth, development, and yield, and provoke the reconfiguration of metabolic processes and activation of key stress responses. However, a remaining question is the performance of pennycress under waterlogging in field environments, particularly in response to heavy spring precipitation, but also to repeated flood events throughout the growing season. Additionally, pennycress fields are susceptible to standing water following snow melt during the winter, leading to partial or full submergence of rosettes. The impact of flooding stress on pennycress phenology, growth, yield, and oil quality in field environments at rosette and reproductive stages should be further evaluated. Moreover, the addition of metabolomic analyses of waterlogged pennycress will elucidate and/or provide additional insight into the results reported here, particularly in relation to the utilization of carbohydrate reserves for survival. Lastly, a unique aspect of this study was the inclusion of seeds and silicle tissue for fatty-acid analysis and RNA-sequencing, where we found decreased levels of VLCFAs and significant enrichment of downregulated genes involved in fatty acid biosynthesis in MN106 under waterlogging. With the seeds being the marketable portion of pennycress, it is crucial to investigate how unfavorable conditions such as waterlogging impact seed development and quality.

As the tools and resources for the pennycress community continue to expand, the analysis of naturally diverse pennycress accessions will be valuable for exploring natural variation in waterlogging tolerance at the morphological and molecular level. While variation in yield was observed between the reference lines (MN106 and SP32-10) under waterlogging, tapping into a larger pool of genetic diversity will help elucidate complex molecular mechanisms involved in flooding tolerance. Furthermore, functional validation of candidate genes through EMS mutagenesis is made possible by an already curated EMS mutant library (Chopra et al., 2018). Additionally, the protocols for gene-editing are well-established in pennycress (McGinn et al., 2019). Functional validation of candidate genes involved in waterlogging tolerance will equip breeders with new knowledge for the development of these resilient lines. As a cash crop in the early stages of commercialization, the quick development and incorporation of climate-resilient pennycress will result in a more robust and successful crop.

## Data availability statement

The RNA-seq data for this project is available at the NCBI BioProject PRJNA1135492 or by request. All supplementary data files may be found at https://github.com/combsgiroir/combsgiroir and custom codes used for data analysis may be found at https://github.com/combsgiroir/Code-for-RNA-seq_Analysis.

## Author contributions

RCG and ARG conceptualized the project. RCG, MEP, and WBP developed methodology and performed experiments. RCG curated data. RCG, MS, HC, MM, EP, AT, and DAJ developed bioinformatics methodology, performed data analyses, and data visualization. RCG, MS, HC, MM, EP, AT, DAJ, ARG contributed to data interpretation. RCG, MS, HC, MM, EP, AT, MEP, WBP, DAJ, and ARG wrote the original draft. RCG and ARG reviewed and edited the manuscript. ARG supervised the project.

## Funding

This research was funded by the U.S. Department of Energy, Office of Science, Office of Biological and Environmental Research, Genomic Science Program [DE-SC0021286]. Combs-Giroir was also financially supported in part by the National Science Foundation Graduate Research Fellowship Program.

## Conflict of interest

The authors declare that the research was conducted in the absence of any commercial or financial relationships that could be construed as a potential conflict of interest.

## Acknowledgements

We would like to thank Ben Phillips, Rosemary Ball, Annabel Shim, Katie Fulcher, and Alex Koopmans for their assistance with data collection and plant maintenance, as well as Cullen Dixon for assistance with tissue collections and RNA-seq data analysis. We would like to thank the Ohio Supercomputer Center for use of computing resources throughout this project. We would also like to thank Jack McCoy for his instruction on the LICOR-6800 instrument and assistance with physiological data collection. Additionally, we would like to thank Gary Posey for maintenance and assistance with greenhouse/growth chamber spaces. Lastly, we would like to thank the Integrated Pennycress Resilience Project (IPReP) team for their insight and constructive feedback related to our project and everything pennycress.

Pathways shown in Figure 6 were created with BioRender.com.

## Notes

https://github.com/combsgiroir/combsgiroir

https://github.com/combsgiroir/Code-for-RNA-seq_Analysis

### Competing Interest Statement

The authors have declared no competing interest.

