## Supplemental Figures for "Morpho-physiological and transcriptomic responses of field pennycress to waterlogging"

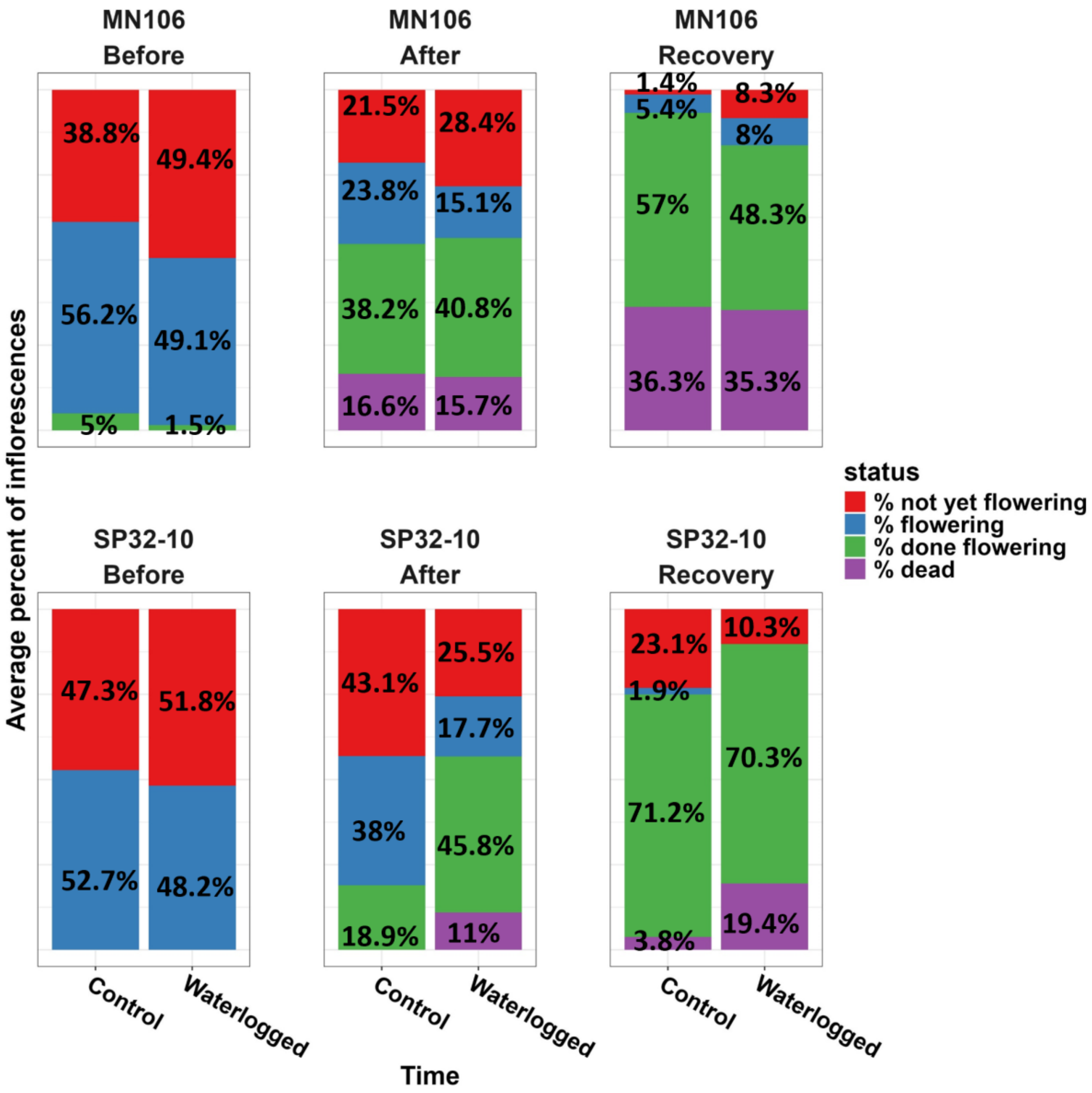


Supplemental Figure 1. Stacked bar plots representing the mean status of inflorescences immediately before waterlogging, immediately after the 7d waterlogging treatments, and after 1 week of recovery from the 7d waterlogging treatments of MN106 and SP32-10 in the growth chamber experiment. Percentages were based on the status of inflorescences divided by the total number of inflorescences per plant. Sample size = 6.

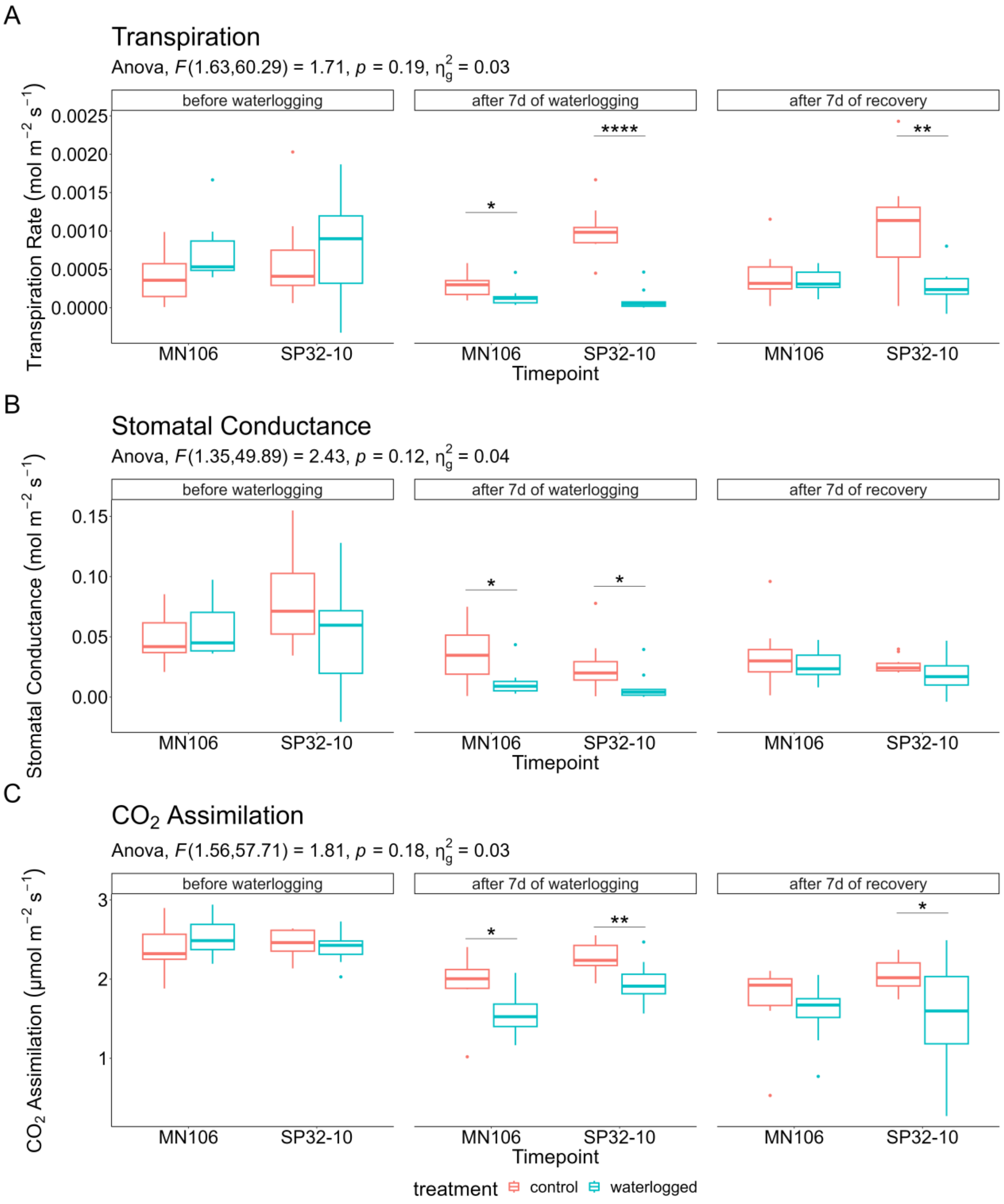


Supplemental Figure 2. Physiological responses before waterlogging, after 7 days of waterlogging, and after 7 days of recovery from waterlogging in the greenhouse experiment. A) Transpiration rate (log transformed), B) Stomatal conductance (log transformed), C) $CO_2$ assimilation (log transformed). * = p-value < 0.05, ** = p-value < 0.01, *** = p-value < 0.001, **** = p-value < 0.0001. Subtitle represents three-way mixed ANOVA result with F statistic, p-value, and generalized eta-squared.

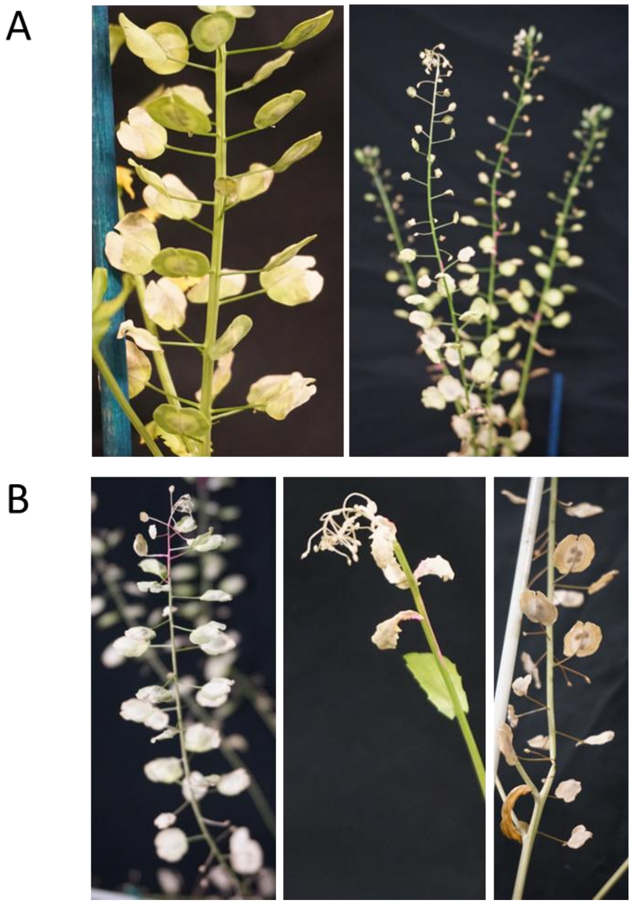


Supplemental Figure 3. Signs of early senescence on silicles and inflorescences after 1 week of recovery from waterlogging in the growth chamber experiment in A) MN106 and B) SP32-10.

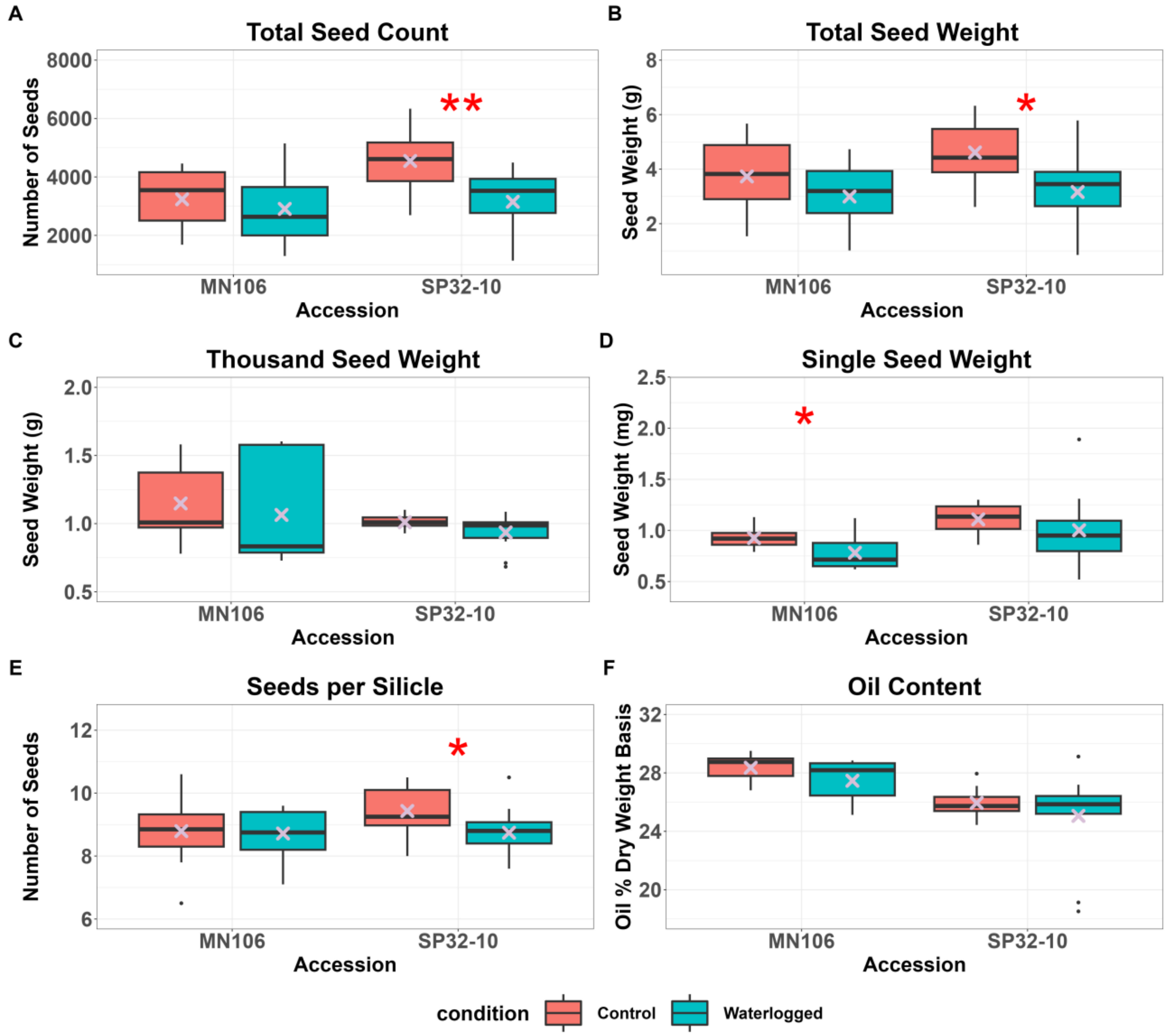


Supplemental Figure 4. Seed yield at the time of harvest following the waterlogging treatment in the greenhouse experiment. A) Total seed count, B) Total seed weight, C) Thousand seed weight, D) Single seed weight, E) Number of seeds per silicle, and F) Total seed oil content. * = p-value < 0.05, ** = p-value < 0.01, ***=p-value < 0.001, “X symbol” represents the mean.

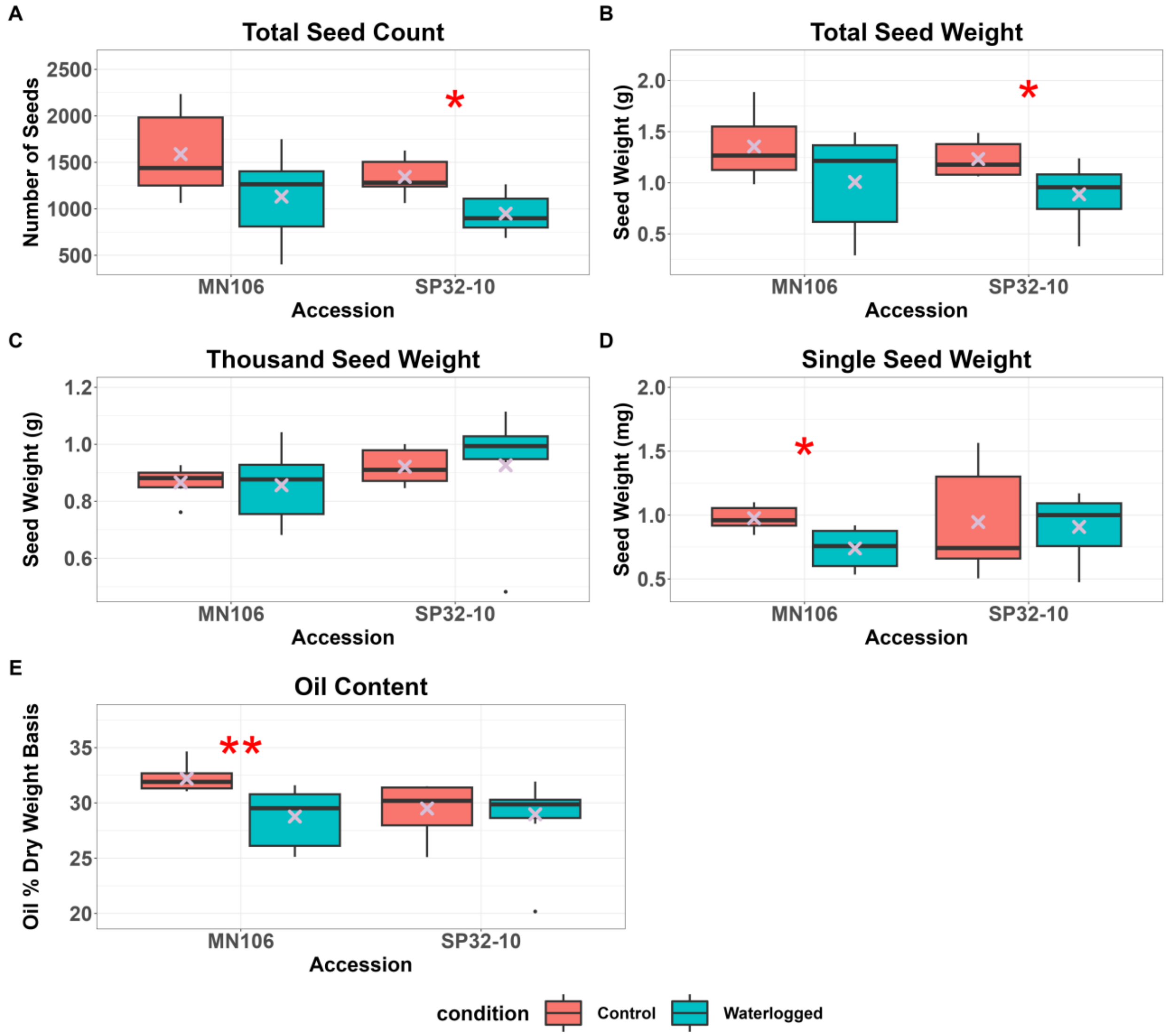


Supplemental Figure 5. Seed yield at the time of harvest following the waterlogging treatment in the growth chamber experiment. A) Total seed count, B) Total seed weight, C) Thousand seed weight, D) Single seed weight, and E) Oil content. * = p-value < 0.05, ** = p-value < 0.01, ***=p-value < 0.001, "X symbol" represents the mean.

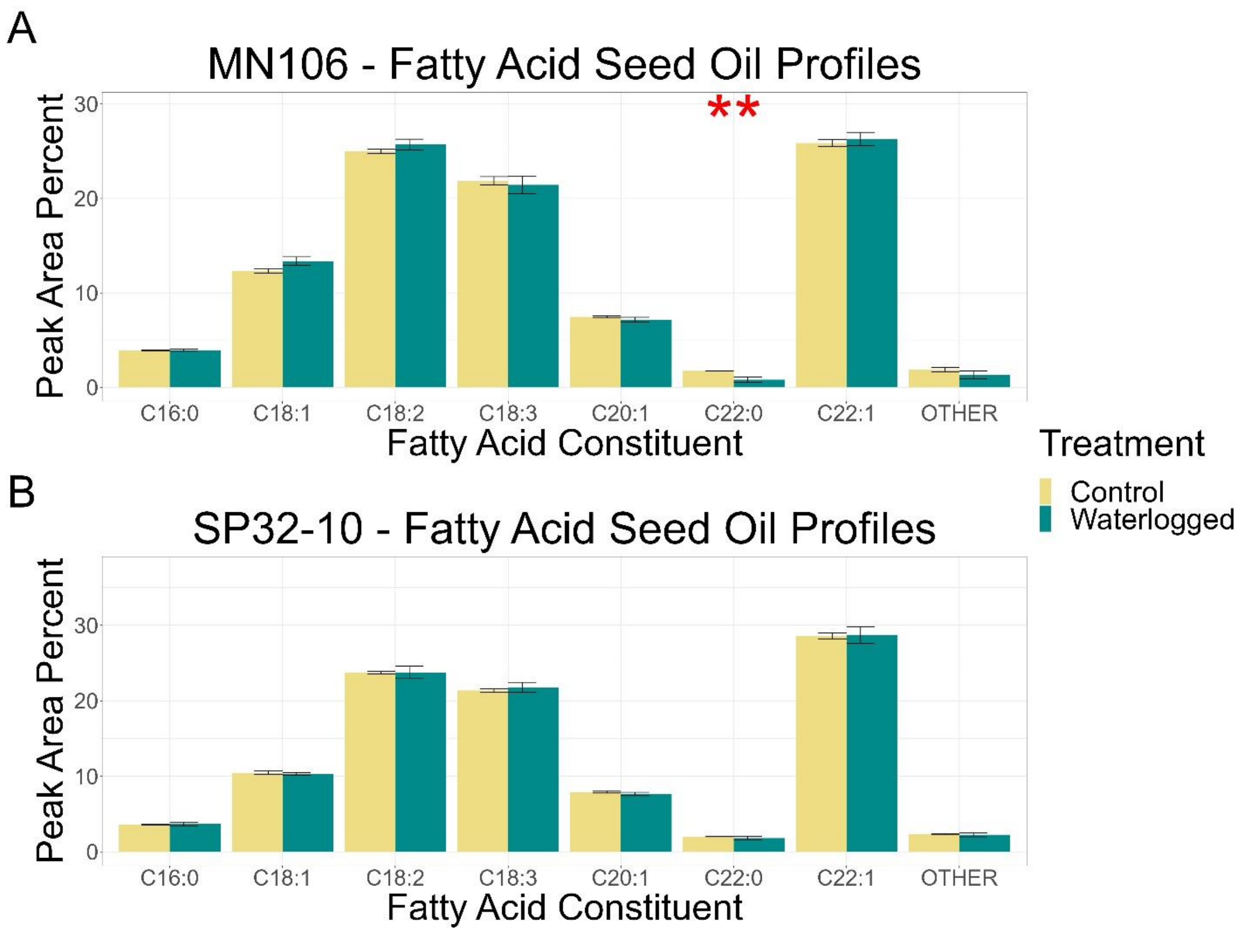


Supplemental Figure 6. Fatty acid oil profiles of mature seed after waterlogging treatment in the growth chamber experiment in A) MN106 and B) SP32-10. Fatty acid constituents are C16:0 = palmitic acid, C18:1 = oleic acid, C18:2 = linoleic acid, C18:3 = linolenic acid, C20:1 = eicosenoic acid, C22:0 = behenic acid, C22:1 = erucic acid. Level of significance reported from Welch's Two Sample t-test. * = p-value < 0.05, ** = p-value < 0.01, ***=p-value < 0.001, bars denote standard error.

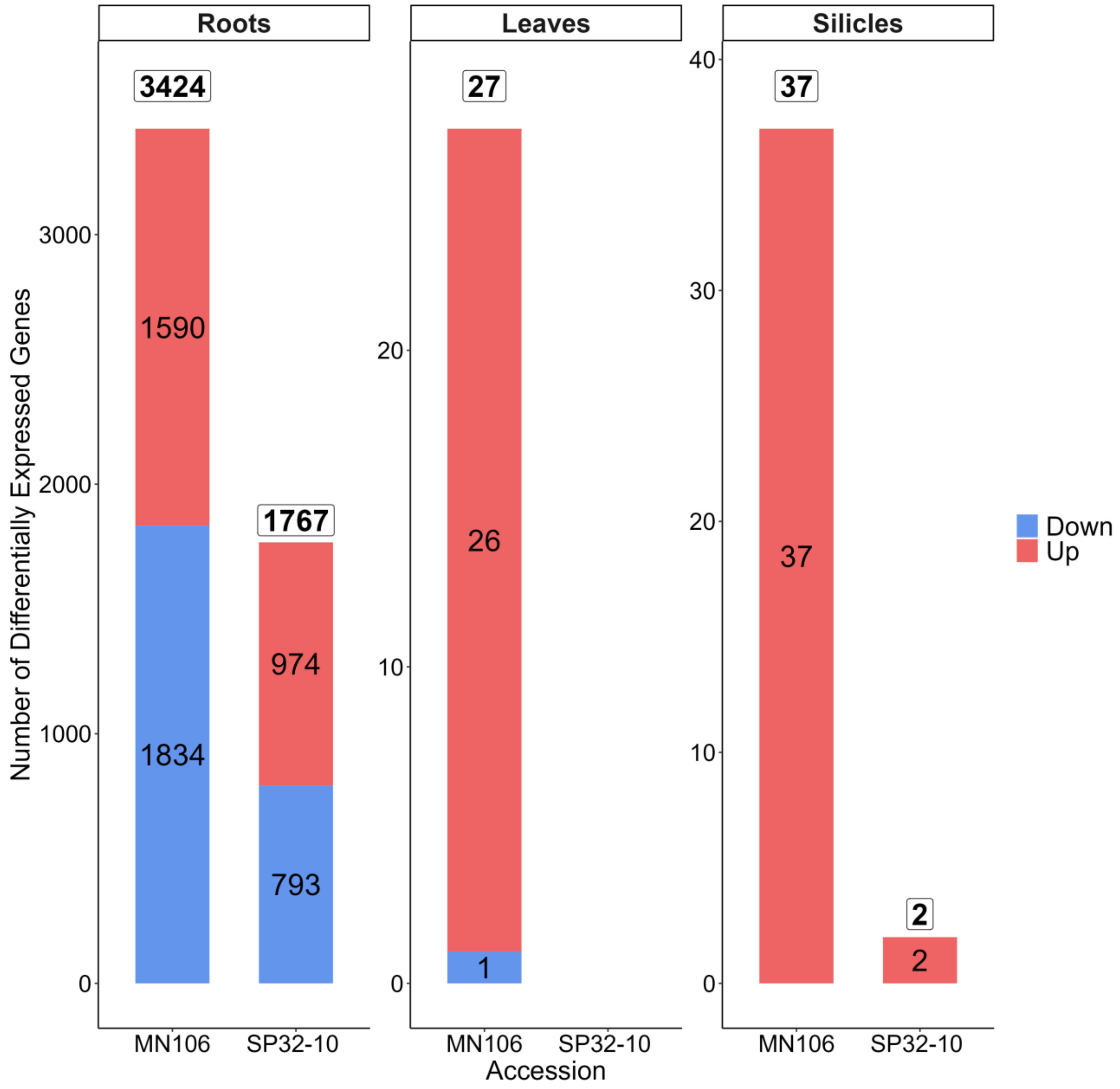


Supplemental Figure 7. Stacked bar plot representing the number of up and downregulated differentially expressed genes in roots, leaves, and silicles of MN106 and SP32-10 waterlogged compared to control samples.

Supplemental Figure 8. See attached file.
A dendrogram of the MENTOR output showing clusters of genes ordered by their functional or mechanistic relationships, along with a heatmap indicating upregulation (red) or downregulation (blue) in waterlogged vs control root samples. Clades with red boxes were chosen for discussion in the text.

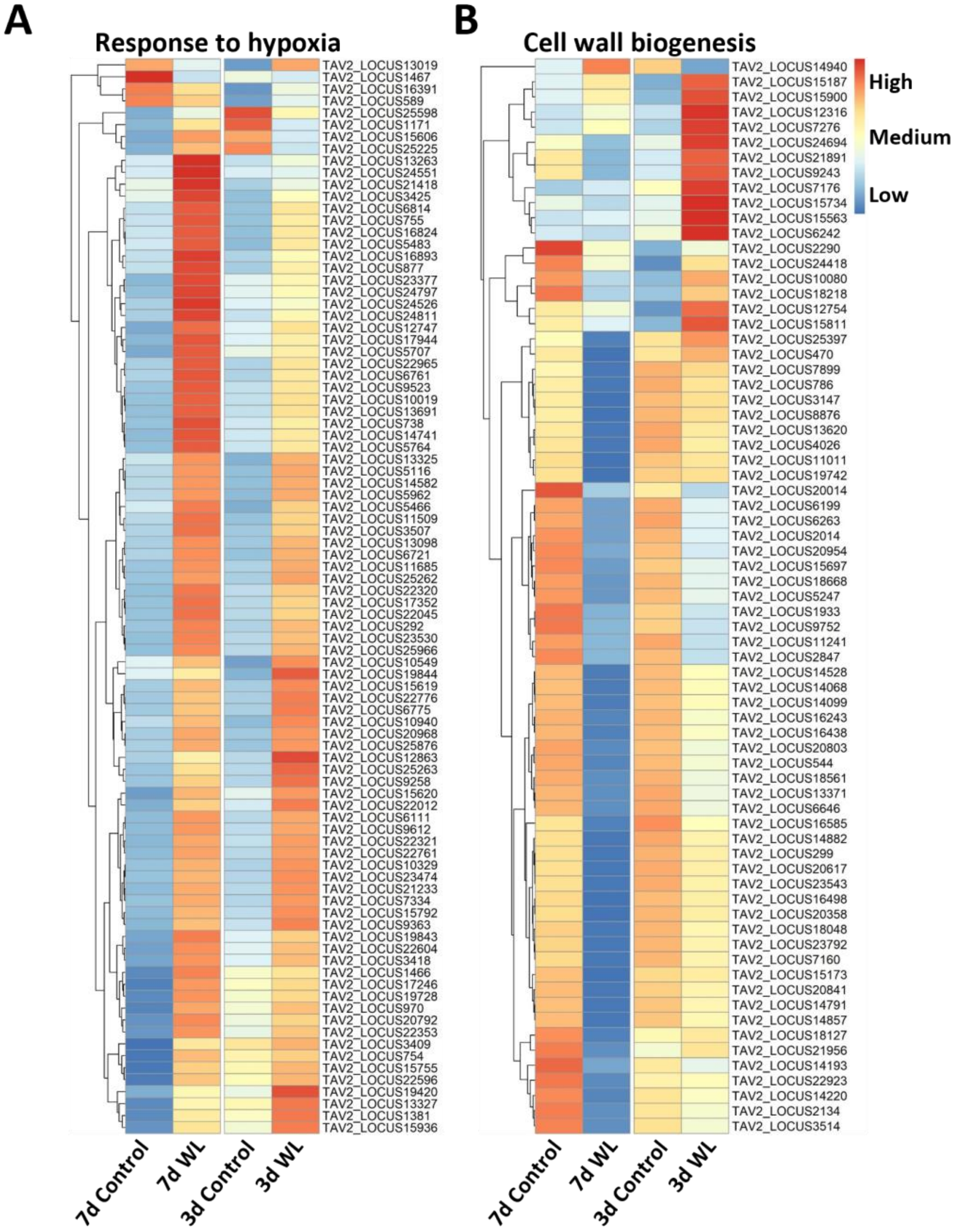


Supplemental Figure 9. Heatmaps showing root gene expression of 7 days and 3 days of waterlogging with controls. A) Gene expression of genes falling under response to hypoxia enriched GO term, and B) Gene expression of genes falling under cell wall biogenesis enriched GO term. The heatmap scale was applied by row or gene.

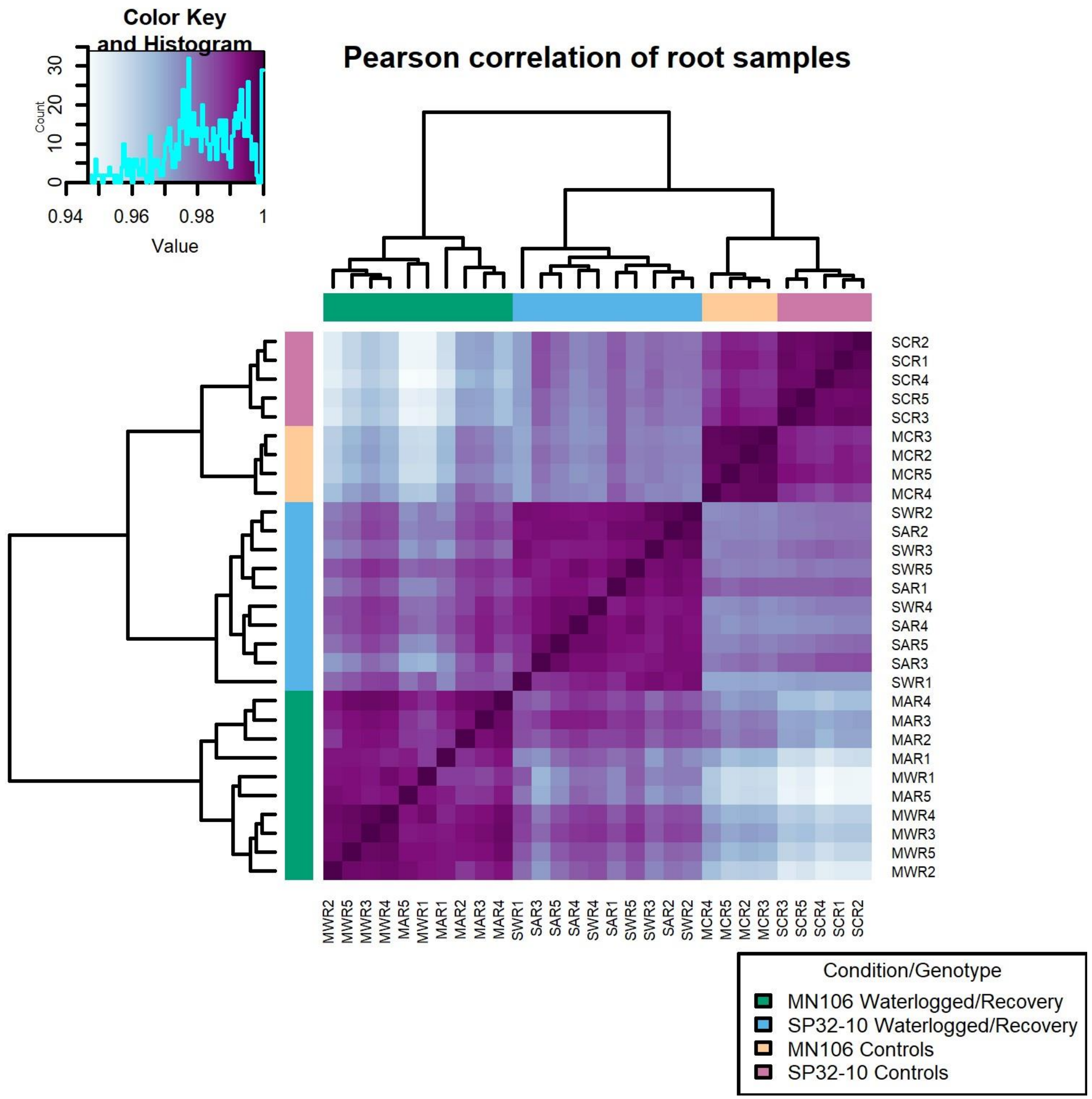


Supplemental Figure 10. Pearson correlation of all root samples with normalized counts as input. Biological replicates MWR1-5 and SWR1-5 indicate MN106 and SP32-10 waterlogged root samples, whereas MAR1-5 and SAR1-5 indicate recovery root samples. MCR1-5 and SCR 1-5 are control root samples.

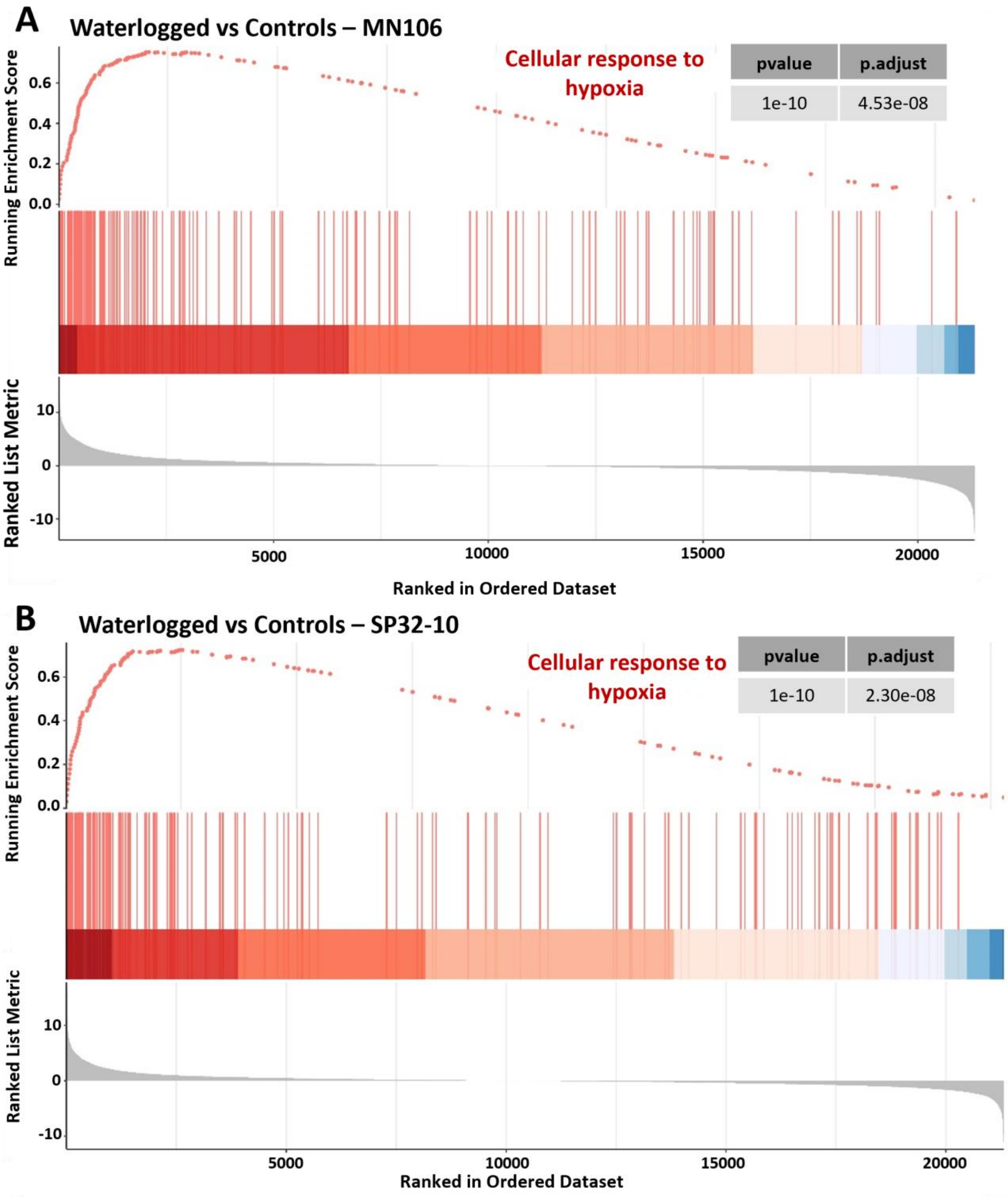
A
Waterlogged vs Controls – MN106
Cellular response to hypoxia
pvalue
p.adjust
1e-10
4.53e-08
Running Enrichment Score
0.0
0.2
0.4
0.6
Ranked List Metric
10
0
-10
5000
10000
15000
20000
Ranked in Ordered Dataset
B
Waterlogged vs Controls – SP32-10
Cellular response to hypoxia
pvalue
p.adjust
1e-10
2.30e-08
Running Enrichment Score
0.0
0.2
0.4
0.6
Ranked List Metric
10
0
-10
5000
10000
15000
20000
Ranked in Ordered Dataset

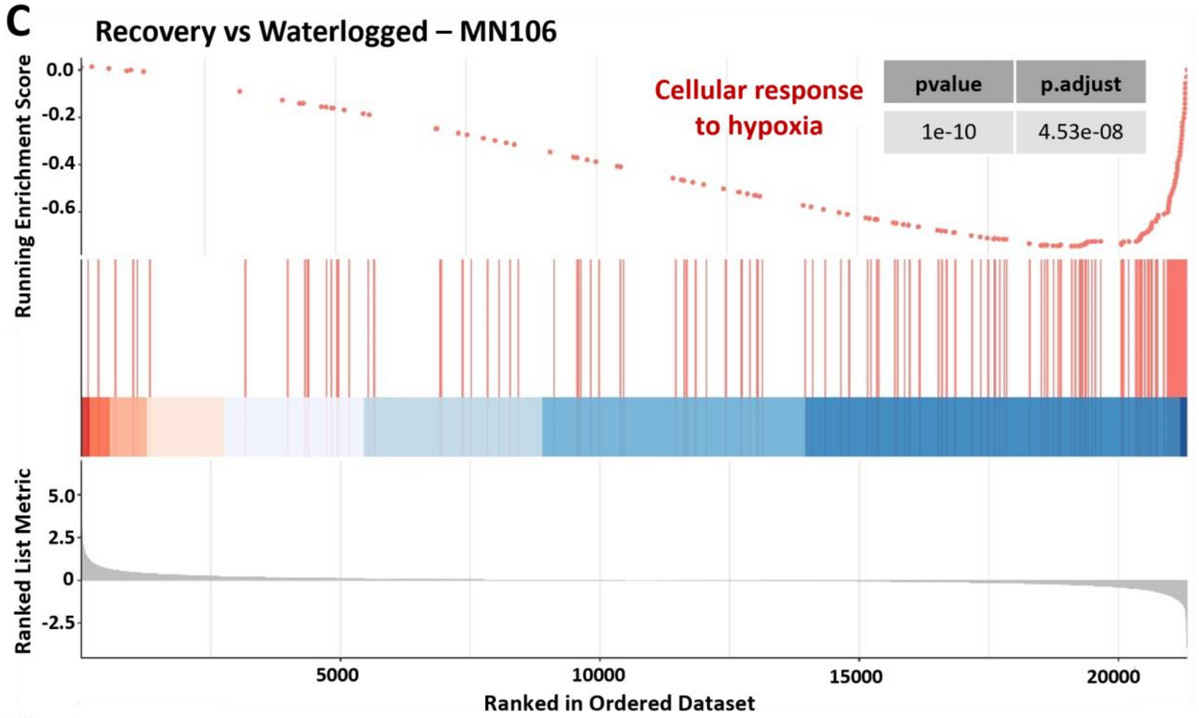
C
Recovery vs Waterlogged – MN106
Cellular response to hypoxia
pvalue
p.adjust
1e-10
4.53e-08
Running Enrichment Score
0.0
-0.2
-0.4
-0.6
Ranked List Metric
5.0
2.5
0
-2.5
5000
10000
15000
20000
Ranked in Ordered Dataset

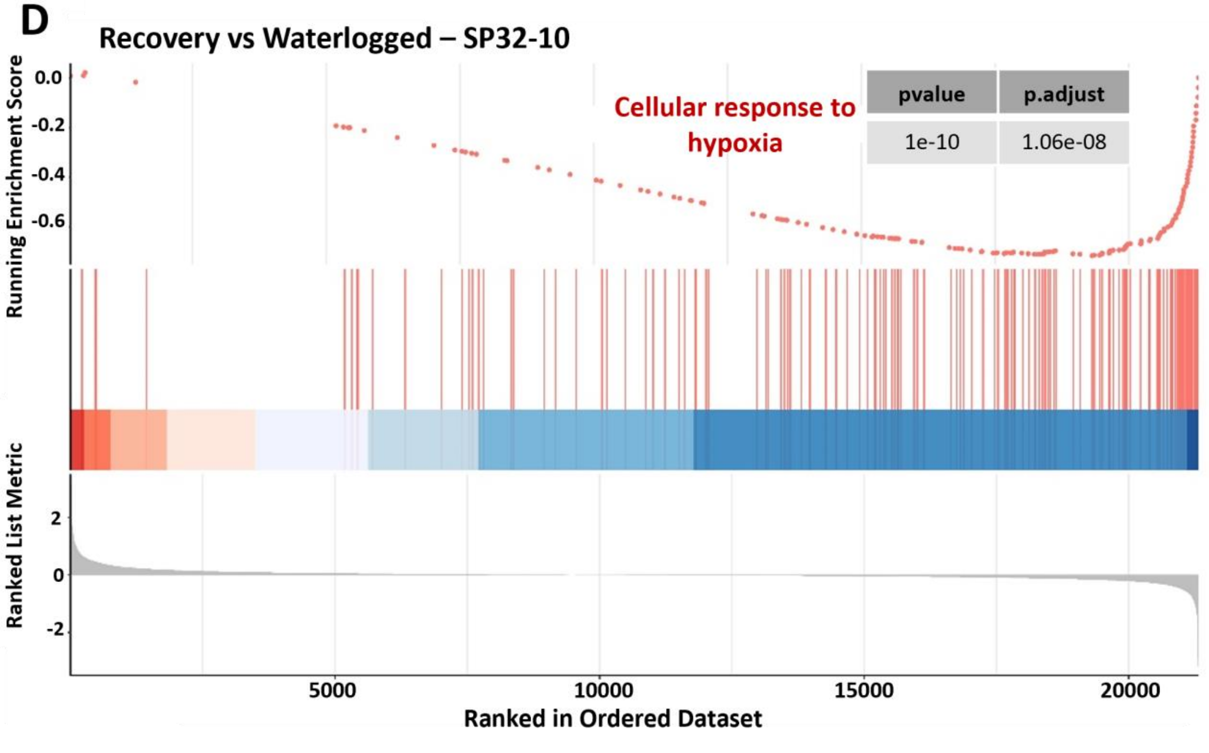
D
Recovery vs Waterlogged – SP32-10
Cellular response to hypoxia
pvalue
p.adjust
1e-10
1.06e-08
Running Enrichment Score
0.0
-0.2
-0.4
-0.6
Ranked List Metric
2
0
-2
5000
10000
15000
20000
Ranked in Ordered Dataset

E

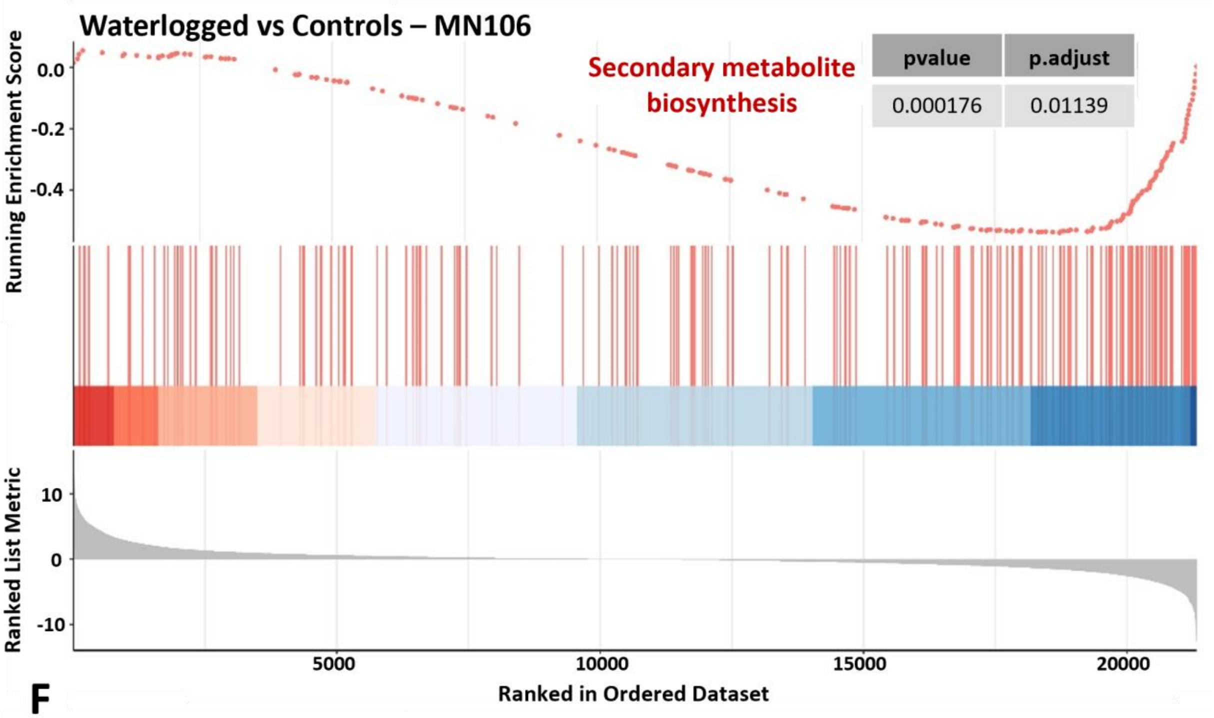

Waterlogged vs Controls – MN106
Secondary metabolite biosynthesis
pvalue
p.adjust
0.000176
0.01139
Running Enrichment Score
0.0
-0.2
-0.4
Ranked List Metric
10
0
-10
5000
10000
15000
20000
Ranked in Ordered Dataset


F

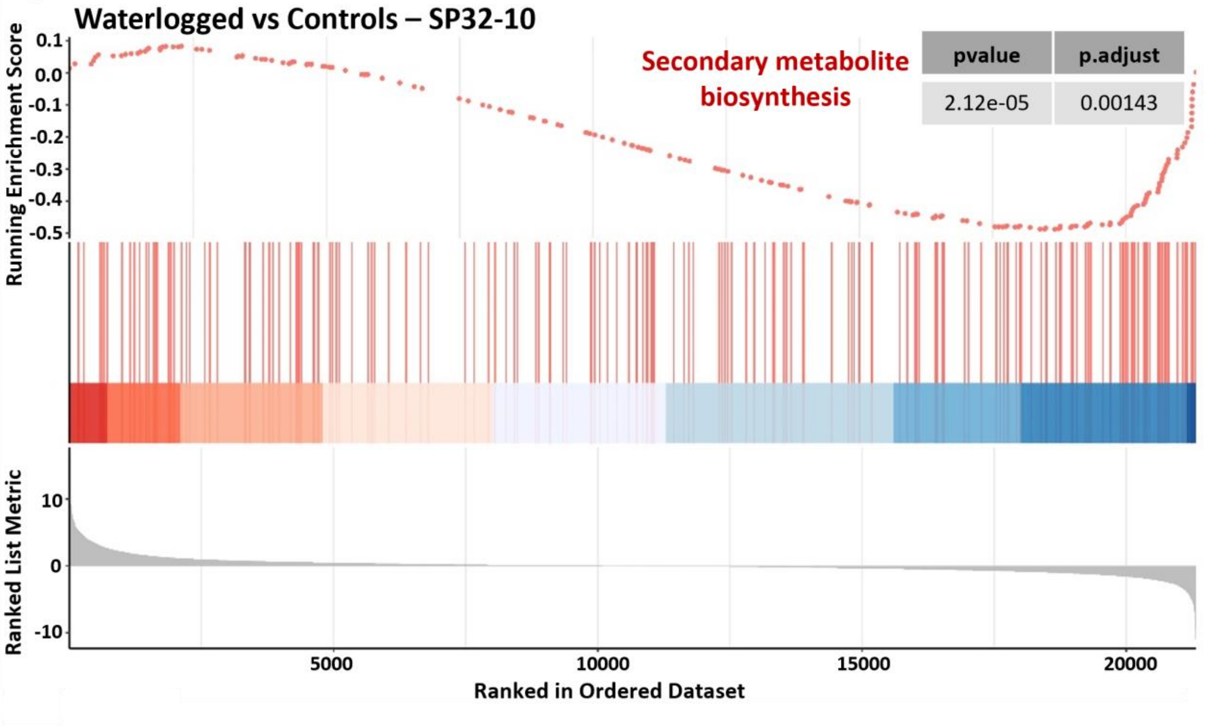

Waterlogged vs Controls – SP32-10
Secondary metabolite biosynthesis
pvalue
p.adjust
2.12e-05
0.00143
Running Enrichment Score
0.1
0.0
-0.1
-0.2
-0.3
-0.4
-0.5
Ranked List Metric
10
0
-10
5000
10000
15000
20000
Ranked in Ordered Dataset

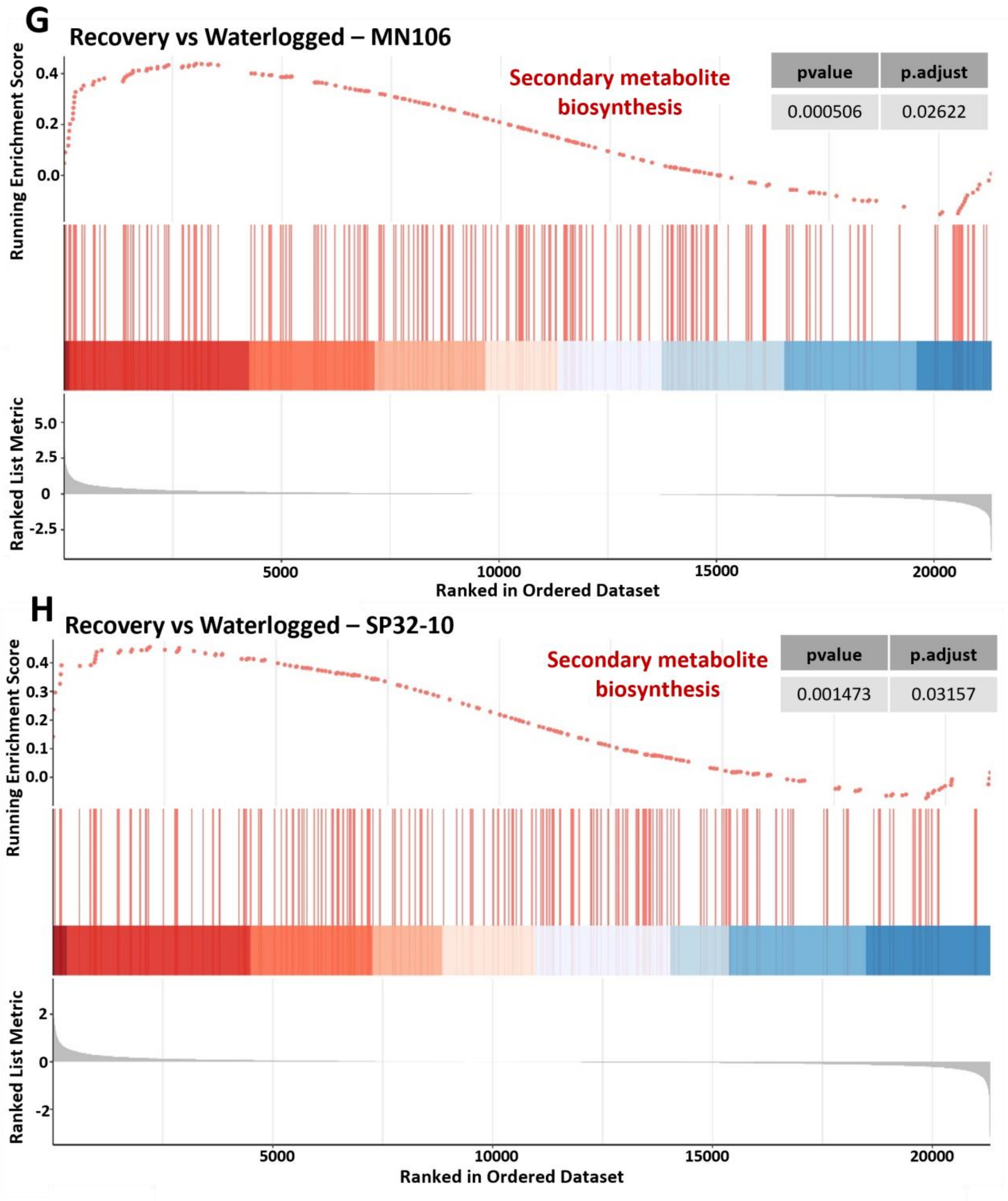


Supplemental Figure 11. Running score and pre-ranked list plot of three hours of recovery compared to 7 days of waterlogging in roots. The enrichment score represents the degree to which a gene set is over-represented at the top or bottom of the ranked list. The ranked list was ordered by a decreasing log2FC. Each gene in the enriched category has a dot for the enrichment score and a vertical line for its position in the ranked list. A-B) Genes involved in hypoxia in waterlogged vs control roots, C-D) Genes involved in hypoxia in recovery vs waterlogged roots (MN-REC vs MN-WL and SP-REC vs SP-WL), E-F) Genes involved in secondary metabolite biosynthesis in waterlogged vs control roots, G-H) Genes involved in secondary metabolite biosynthesis in recovery vs waterlogged roots (MN-REC vs MN-WL and SP-REC vs SP-WL)
