## Supplemental Tables for "Morpho-physiological and transcriptomic responses of field pennycress to waterlogging"

Supplemental Table 1. Means and standard deviations of morphological traits of waterlogged and control pennycress immediately after waterlogging in the growth chamber experiment.

| | **MN106** | | | **SP32-10** | | |
|---|---|---|---|---|---|---|
| | **Control** | **Waterlogged** | ***P-value*** | **Control** | **Waterlogged** | ***P-value*** |
| **Height (cm)** | 64.42 ± 4.67$^{ab}$ | 59.75 ± 9.40$^{b}$ | 0.421 | 70.92 ± 3.83$^{a}$ | 73.25 ± 2.27$^{a}$ | 0.235 |
| **Branch #** | 12.33 ± 4.03$^{b}$ | 12.67 ± 6.38$^{b}$ | 0.916 | 21 ± 4.20$^{a}$ | 19.5 ± 3.62$^{ab}$ | 0.523 |
| **Silicle #** | 306.33 ± 67.5$^{a}$ | 271.67 ± 113.4$^{a}$ | 0.538 | 291.5 ± 69.6$^{a}$ | 282 ± 68.4$^{a}$ | 0.816 |
| **Aborted Silicles (%)** | 1.38 ± 1.74$^{b}$ | 3.76 ± 4.60$^{b}$ | 0.277 | 6.99 ± 3.08$^{ab}$ | 13.54 ± 8.81$^{a}$ | 0.135 |

Sample size = 6. P-values derived from Welch’s t-test between treatments for each accession. Groups with the same letter next to the standard deviation are not statistically different based on Tukey’s HSD test following a two-way ANOVA between accession and treatment.

Supplemental Table 2. Mean values and standard deviations of the status of inflorescences before waterlogging, after waterlogging, and after 1 week of recovery in MN106 and SP32-10 in the growth chamber experiment.

| | MN106 | | | SP32-10 | | |
|---|---|---|---|---|---|---|
| | **Control** | **Waterlogged** | ***P-value*** | **Control** | **Waterlogged** | ***P-value*** |
| *Before Waterlogging* | | | | | | |
| not yet flowering | 0.39 ± 0.11[a] | 0.49 ± 0.15[a] | 0.199 | 0.47 ± 0.14[a] | 0.52 ± 0.11[a] | 0.546 |
| flowering | 0.56 ± 0.17[a] | 0.49 ± 0.17[a] | 0.487 | 0.53 ± 0.14[a] | 0.48 ± 0.11[a] | 0.546 |
| done flowering | 0.05 ± 0.12[a] | 0.02 ± 0.04[a] | 0.523 | 0[a] | 0[a] | NA |
| dead | 0 | 0 | NA | 0 | 0 | NA |
| *After Waterlogging* | | | | | | |
| not yet flowering | 0.21 ± 0.11[a] | 0.28 ± 0.22[a] | 0.507 | 0.43 ± 0.08[a] | 0.26 ± 0.18[a] | 0.069 |
| flowering | 0.24 ± 0.24[a] | 0.15 ± 0.22[a] | 0.529 | 0.38 ± 0.18[a] | 0.18 ± 0.15[a] | 0.059 |
| done flowering | 0.38 ± 0.24[a] | 0.41 ± 0.12[a] | 0.821 | 0.19 ± 0.23[a] | 0.46 ± 0.22[a] | 0.063 |
| dead | 0.17 ± 0.12[a] | 0.16 ± 0.18[a] | 0.924 | 0[a] | 0.11 ± 0.09[a] | 0.023* |
| *1 Week of Recovery* | | | | | | |
| not yet flowering | 0.01 ± 0.03[a] | 0.08 ± 0.20[a] | 0.447 | 0.23 ± 0.20[a] | 0.10 ± 0.08[a] | 0.183 |
| flowering | 0.05 ± 0.09[a] | 0.08 ± 0.11[a] | 0.668 | 0.02 ± 0.03[a] | 0[a] | 0.211 |
| done flowering | 0.57 ± 0.13[a] | 0.48 ± 0.13[a] | 0.269 | 0.71 ± 0.18[a] | 0.70 ± 0.13[a] | 0.925 |
| dead | 0.36 ± 0.15[a] | 0.35 ± 0.23[a] | 0.936 | 0.04 ± 0.09[b] | 0.19 ± 0.17[ab] | 0.083 |

Means were determined by the number of inflorescences for each status (not yet flowering, flowering, done flowering, dead) divided by the total number of inflorescences on the plant. Sample size = 6. Asterisk denotes statistical significance of < 0.05. Groups with the same letter next to the standard deviation are not statistically different based on Tukey's HSD test following a two-way ANOVA between accession and treatment for each time point.

Supplemental Table 3. Means and standard deviations of morphological traits of waterlogged and control pennycress after 1 and 2 weeks of recovery from waterlogging in the growth chamber experiment.

| | **MN106** | | | **SP32-10** | | |
|---|---|---|---|---|---|---|
| | **Control** | **Waterlogged** | ***P-value*** | **Control** | **Waterlogged** | ***P-value*** |
| *1 week* | | | | | | |
| **Height (cm)** | 64.4 ± 4.67[bc] | 60.5 ± 9.22[c] | 0.572 | 71.4 ± 4.36[ab] | 74.5 ± 3.45[a] | 0.206 |
| **Branch #** | 12.3 ± 4.03[b] | 13.5 ± 5.72[b] | 0.693 | 24.5 ± 7.09[a] | 20.8 ± 3.82[ab] | 0.298 |
| **Silicle #** | 335.2 ± 73.5[a] | 282.2 ± 111.8[a] | 0.358 | 374.3 ± 58.0[a] | 326.3 ± 57.8[a] | 0.182 |
| **Senesced Silicles (%)** | 0[a] | 1.70 ± 3.9[a] | 0.176 | 0[a] | 18.8 ± 30.7[a] | 0.010** |
| **Aborted Silicles (%)** | 2.76 ± 3.24[b] | 5.71 ± 6.85[b] | 0.371 | 21.8 ± 7.82[a] | 21.5 ± 6.19[a] | 0.945 |
| *2 weeks* | | | | | | |
| **Silicle #** | 336.3 ± 74.9[a] | 291.5 ± 121.4[a] | 0.463 | 392.7 ± 78.7[a] | 333.3 ± 62.3[a] | 0.394 |
| **Senesced Silicles (%)** | 4.53 ± 6.0[a] | 24.7 ± 25.3[a] | 0.111 | 7.03 ± 5.44[a] | 30.1 ± 34.4[a] | 0.009** |

Sample size = 6. Asterisk denotes statistical significance of < 0.05. P-values derived from Welch's t-test between treatments for each accession. Groups with the same letter next to the standard deviation are not statistically different based on Tukey's HSD test following a two-way ANOVA between accession and treatment.

Supplemental Table 4. Means and standard deviations of morphological traits of waterlogged and control plants at the time of harvest in the growth chamber experiment.

| | MN106 | | | SP32-10 | | |
|---|---|---|---|---|---|---|
| | **Control** | **Waterlogged** | ***P-value*** | **Control** | **Waterlogged** | ***P-value*** |
| **Height (cm)** | 64.8 ± 4.75[ab] | 59.8 ± 10.2[b] | 0.465 | 71.4 ± 4.36[a] | 74.5 ± 3.45[a] | 0.206 |
| **Reproductive Height (cm)** | 30.3 ± 4.34[a] | 27.5 ± 5.14[a] | 0.341 | 31.3 ± 4.37[a] | 33 ± 3.29[a] | 0.474 |
| **Primary Branch #** | 5.17 ± 1.33[ab] | 4.5 ± 2.17[b] | 0.538 | 7.83 ± 1.94[a] | 7 ± 2.53[ab] | 0.540 |
| **Branch #** | 12.3 ± 4.03[b] | 13.5 ± 5.72[b] | 0.693 | 24.5 ± 7.09[a] | 20.8 ± 3.82[ab] | 0.298 |
| **Silicle #** | 339.3 ± 74.9[a] | 306.3 ± 133.6[a] | 0.612 | 401.7 ± 72.3[a] | 352.8 ± 61.1[a] | 0.394 |
| **Aborted Silicles (%)** | 16.9 ± 3.1[b] | 25.1 ± 9.39[b] | 0.091 | 35.5 ± 6.17[a] | 44.5 ± 3.23[a] | 0.014* |
| **Maturity** | 45.7 ± 1.86[a] | 44.2 ± 4.02[a] | 0.434 | 42.7 ± 1.03[a] | 41.2 ± 4.49[a] | 0.458 |
| **Shoot Dry Weight (g)** | 4.27 ± 1.05[a] | 3.3 ± 1.20[a] | 0.169 | 3.59 ± 0.46[a] | 3.09 ± 0.73[a] | 0.299 |
| **Total Seed Count** | 1588.2 ± 486.3[a] | 1131.2 ± 500.2[ab] | 0.140 | 1342.8 ± 217.5[ab] | 948.7 ± 223.9[b] | 0.011* |
| **Total Seed Weight (g)** | 1.35 ± 0.34[a] | 1.01 ± 0.51[a] | 0.201 | 1.23 ± 0.19[a] | 0.89 ± 0.31[a] | 0.048* |
| **Thousand Seed Weight (g)** | 0.87 ± 0.06[a] | 0.86 ± 0.14[a] | 0.876 | 0.92 ± 0.07[a] | 0.93 ± 0.23[a] | 0.310 |
| **Single Seed Weight (m g)** | 0.98 ± 0.10[a] | 0.74 ± 0.17[a] | 0.018* | 0.95 ± 0.46[a] | 0.91 ± 0.27[a] | 0.864 |
| **Oil Content (% DWB)** | 32.2 ± 1.12[a] | 28.8 ± 2.60[b] | 0.004** | 29.5 ± 2.21[ab] | 29 ± 3.31[b] | 0.675 |

Sample size = 6. Asterisk denotes statistical significance of < 0.05. DWB = dry weight basis. P-values derived from Welch's t-test between treatments for each accession. Groups with the same letter next to the standard deviation are not statistically different based on Tukey's HSD test following a two-way ANOVA between accession and treatment.

Supplemental Table 5. List of top significant differentially expressed genes (DEGs) in waterlogged vs control roots for MN106 and SP32-10 with log2FC, significance level (padj), and differential expression rank among the total DEG list.

| | | | | MN106 (MN-7WL) | | | SP32-10 (SP-7WL) | | |
|---|---|---|---|---|---|---|---|---|---|
| Pennycress ID | Arabidopsis ID | Gene Name/Putative Function | Abbr. | log2FC | padj | rank | log2FC | padj | rank |
| TAV2_LOCUS11649 | –[A] | *cytochrome P450* | – | 11.53 | 3E-107 | **1** | 7.872 | 1.3E-64 | **3** |
| TAV2_LOCUS281 | AT4G10270 | *WOUND-INDUCED POLYPEPTIDE 4* | *WIP4* | 7.379 | 3.1E-73 | **2** | 7.087 | 3.3E-70 | **2** |
| TAV2_LOCUS6929 | AT2G02010 | *GLUTAMATE DECARBOXYLASE 4* | *GAD4* | 7.561 | 3.7E-64 | **3** | 5.339 | 1E-31 | 30 |
| TAV2_LOCUS17926 | AT1G67100 | *LOB DOMAIN-CONTAINING PROTEIN 40* | *LBD40* | 9.802 | 3.7E-64 | **4** | 6.581 | 4.9E-43 | 16 |
| TAV2_LOCUS19745 | AT5G54960 | *PYRUVATE DECARBOXYLASE 2* | *PDC2* | 8.257 | 5.3E-61 | **5** | 8.634 | 2.8E-75 | **1** |
| TAV2_LOCUS5321 | AT1G64600 | *RIBOSOMAL PROTEIN MS22* | *MS22* | -5.765 | 1.8E-58 | **6** | -4.637 | 9.7E-39 | 18 |
| TAV2_LOCUS23530 | AT4G37710 | *VQ MOTIF-CONTAINING PROTEIN 29* | *VQ29* | 7.909 | 5.8E-56 | **7** | 7.198 | 4.9E-48 | 13 |
| TAV2_LOCUS17395 | AT1G73010 | *PHOSPHATE STARVATION-INDUCED GENE 2* | *ATPS2* | 8.691 | 1.6E-54 | **8** | 3.199 | 9.1E-06 | 597 |
| TAV2_LOCUS5907 | AT3G29970 | B12D protein | – | 9.259 | 1.1E-52 | **9** | 9.984 | 1.7E-62 | **4** |
| TAV2_LOCUS5664 | AT5G26710 | Glutamyl/glutaminyl-tRNA synthetase | – | 5.177 | 4.3E-52 | **10** | 4.624 | 4.9E-44 | 15 |
| TAV2_LOCUS10091 | AT2G19590 | *ACC OXIDASE 1* | *ACO1* | 8.984 | 1.6E-41 | 21 | 9.962 | 5.3E-59 | **5** |
| TAV2_LOCUS13607 | AT2G29870 | Aquaporin-like superfamily protein | – | 9.174 | 2.1E-32 | 47 | 10.21 | 4.7E-57 | **6** |
| TAV2_LOCUS18148 | AT1G77120 | *ALCOHOL DEHYDROGENASE 1* | *ADH1* | 5.595 | 9.9E-50 | 11 | 5.641 | 4.7E-57 | **7** |
| TAV2_LOCUS26083 | AT4G20840 | *OLIGOGALACTURONIDE OXIDASE 2* | *OGOX2* | 4.701 | 2E-41 | 22 | 4.92 | 2.7E-52 | **8** |
| TAV2_LOCUS21261 | – | Unknown - protein kinase domain | – | 6.627 | 1.7E-25 | 103 | 9.447 | 7.4E-51 | **9** |
| TAV2_LOCUS10094 | – | *ACC OXIDASE 1* | *ACO1* | 9.209 | 7.3E-36 | 33 | 9.969 | 1.7E-48 | **10** |

[A]Some genes did not have an Arabidopsis ortholog, therefore, the protein sequences were BLASTed to find a putative function based on conserved domains and orthology to genes from other species.
