## Supplementary figures and images for "Morpho-physiological and transcriptomic responses of field pennycress to waterlogging"

### Supplemental Figure 8

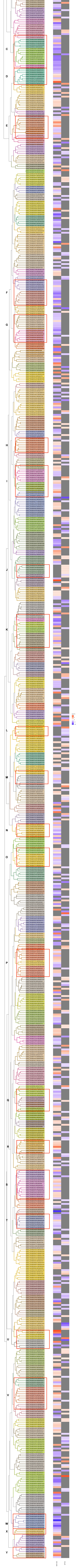
